# Phase Separation-based Antiviral Decoy Particles as Basis for Programmable Broad-spectrum Therapeutics

**DOI:** 10.1101/2024.08.28.610020

**Authors:** Or Willinger, Naor Granik, Sarah Goldberg, Roee Amit

## Abstract

To gain access to cells, viruses employ host proteins as receptors. In soluble form, these receptors are used as decoys to inhibit infection. We fused candidate soluble receptors to an RNA-binding protein, and using synthetic long non-coding RNA (slncRNA) cassettes that can undergo phase-separation we scaffolded the receptor fusions to generate antiviral decoy particles. Using confocal microscopy, we screened antiviral protein candidates by observing changes in phase-separation morphology when incubated with viral-mimicking components. We demonstrated that ACE2 decoy particles bind strongly to the coronavirus RBD, facilitating FRET, while sufficiently sialylated decoy particles form agglutinated structures with RNA peripheries in the presence of a sialolectin. Infection assays show ACE2 decoy particles fully inhibit the Delta and Omicron BA.1 coronavirus variants, and LAMP1 and GYPA decoy particles significantly reduce influenza infection *in-cellulo*. This work establishes a foundation for broad-spectrum antiviral decoy particles, composed of multiple receptors targeting various viruses.

## Main

The COVID-19 pandemic has exposed the world vulnerability to infectious diseases. Current antiviral therapeutic approaches can be roughly divided into two classes. The first includes immunogenic therapies (e.g. vaccines and monoclonal antibodies) that target only a limited number of virus variants at a time, making them narrow spectrum approaches that can rapidly become ineffective as a virus evolves and mutates(1). The second approach includes replication inhibitors, which are either similarly narrow spectrum (e.g. specific protease inhibitors), or can interact with host RNA and DNA polymerases (general nucleic acid analogs) resulting in a wide-array of side effects(1). Consequently, at the present time there is no fully protective antiviral therapeutic strategy to combat viral infectious diseases.

An alternative antiviral therapeutic strategy that has increasingly garnered attention in recent years is the use of soluble receptors. This approach makes use of either the soluble isoforms or a recombinant version of the extracellular domain of host surface proteins to which viruses attach in order begin their infection cycle(2). The soluble-receptor strategy assumes that the virus-host interaction mechanism is an evolutionary bottleneck. This assumption is supported by the observation that no documented virus has naturally changed its target host receptor outside of a lab setting(3). Increasing the local concentration of soluble receptor monomers was shown to be advantageous against SARS-CoV-2(4, 5). However, soluble receptors are inherently unstable protein moieties due to their detachment from their membranal component, leading to a large reduction to their half-life(6), and as a result making them poor therapeutic candidates. To increase stability, soluble receptors are frequently fused to the Fc domain of IgG antibodies(4, 7–9), sometimes even improving the half-maximal inhibitory concentration (IC_50_)(4). However, the use of the Fc domain can trigger antibody-dependent enhancement (ADE), whereby a virus which was bound by an Fc-fused receptor is endocytosed into immune cells, and subsequently triggers inflammatory pathways and cell death(10). As a result, despite their potential to be both a highly specific antiviral therapeutic and one which is insensitive to viral mutations, soluble receptor-based antiviral therapeutics have yet to reach the clinic.

The shortened half-life detriment could be resolved without the use of antibody domains by the phenomena of macromolecular liquid-liquid or liquid-solid phase separation. These phase transitions are mediated by weak molecular interactions, and result in locally dense liquid or gel-like solid within a homogenous solution when a particular concentration threshold of one of the molecular components is reached(11). The process is observed in virtually all cell-types and in many organelles, and has been the subject of many studies over the past decade(12–17). The particles formed in this process are typically characterized as interconnected biomolecular networks that are stabilized by one or more of a set of biophysical interactions, and are described by models such as polyphasic linkage(18), patchy colloid model(14), and others(19–21). Specifically, proteins that tend to phase separate frequently contain domains such as intrinsically disordered domains (IDDs), RNA-binding domains, and various post-translational modifications (PTMs) such as phosphorylation, acetylation, and glycosylation(22, 23). These domains and PTMs serve as points of contact or scaffolds for formation of interconnected molecular networks, and the critical concentration of the phase separation process is in part dependent on the number of points of contact between macromolecules, often defined as their valency(14, 19).

We recently showed that we can create synthetic liquid-gel phase-separated biocondensates termed synthetic RNA-protein (sRNP) granules both in vitro and in-cellulo in various cell types using programable synthetic long non-coding RNA (slncRNA) molecules featuring a cassette of connected hairpin structures, and a matching RNA binding protein (a tandem-dimer PP7-coat protein fused to mCherry [tdPCP-mCherry]) (11, 24). In this work, we increase the complexity of the tdPCP-slncRNA sRNP granules by fusing soluble receptor moieties to the tdPCP, thus endowing the granules with presumptive antiviral functionality. To test for potential antiviral functionality, we quantified changes to granule structure and composition via fluorescence microscopy upon introduction of a third molecular component that mimics the host-binding component of either SARS-CoV-2 or Influenza A. We then confirmed the antiviral functionality for SARS-CoV-2 and influenza A in viral entry assays on the best-performing granule variants. Consequently, we developed a phase-separation-based method to detect and quantify different types of molecular binding events for granule-binding proteins and their ligands. Finally, we employed the same phase-separated granule platform to inhibit two SARS-CoV-2 variants and an influenza A strain.

## Results

### Using phase-separation to confirm functionality of antiSARS2 decoy granules

We hypothesized that sRNP granules can be adapted to form stable antiviral decoy nanoparticles by fusing the tdPCP moiety to the soluble receptor domain of a virus target receptor, and that such particles can be screened for antiviral functionality via changes to the phase-separated granule structure when exposed to the viral receptor binding moiety. To test our strategy (Figure 1A) for developing antiviral granules, we opted to first use our putative antiSARS2 granules, reported previously(25). In brief, we demonstrated that FITC-labelled polystyrene beads coated with the SARS-CoV-2 receptor binding domain (RBD) form tightly bound or agglutinated clusters with sRNP granules, composed of a slncRNA containing eight tdPCP binding sites and tdPCP-mCherry-ACE2 protein (Supplementary Figure S1). To characterize the putative antiSARS2 granules, we employed a dual-label fluorescence microscopy approach with the slncRNA labelled with AF405-Uracil during synthesis, together with the tdPCP-mCherry-ACE2 moiety (Figure 1V). The image shows formation of either protein condensates (red) with faint slncRNA signal, or co-localized protein and slncRNA spots (blue), indicating formation of protein-RNA biocondensates or granules. We then correlated the red and blue intensities of 339 co-localized structures over 27 field-of-views (FoVs) and found a significant correlation between red and blue intensities (Figure 1C and Supplementary Figure S2A), providing support for the formation of granulated structure with a range of protein to slncRNA ratios.

**Figure 1.**
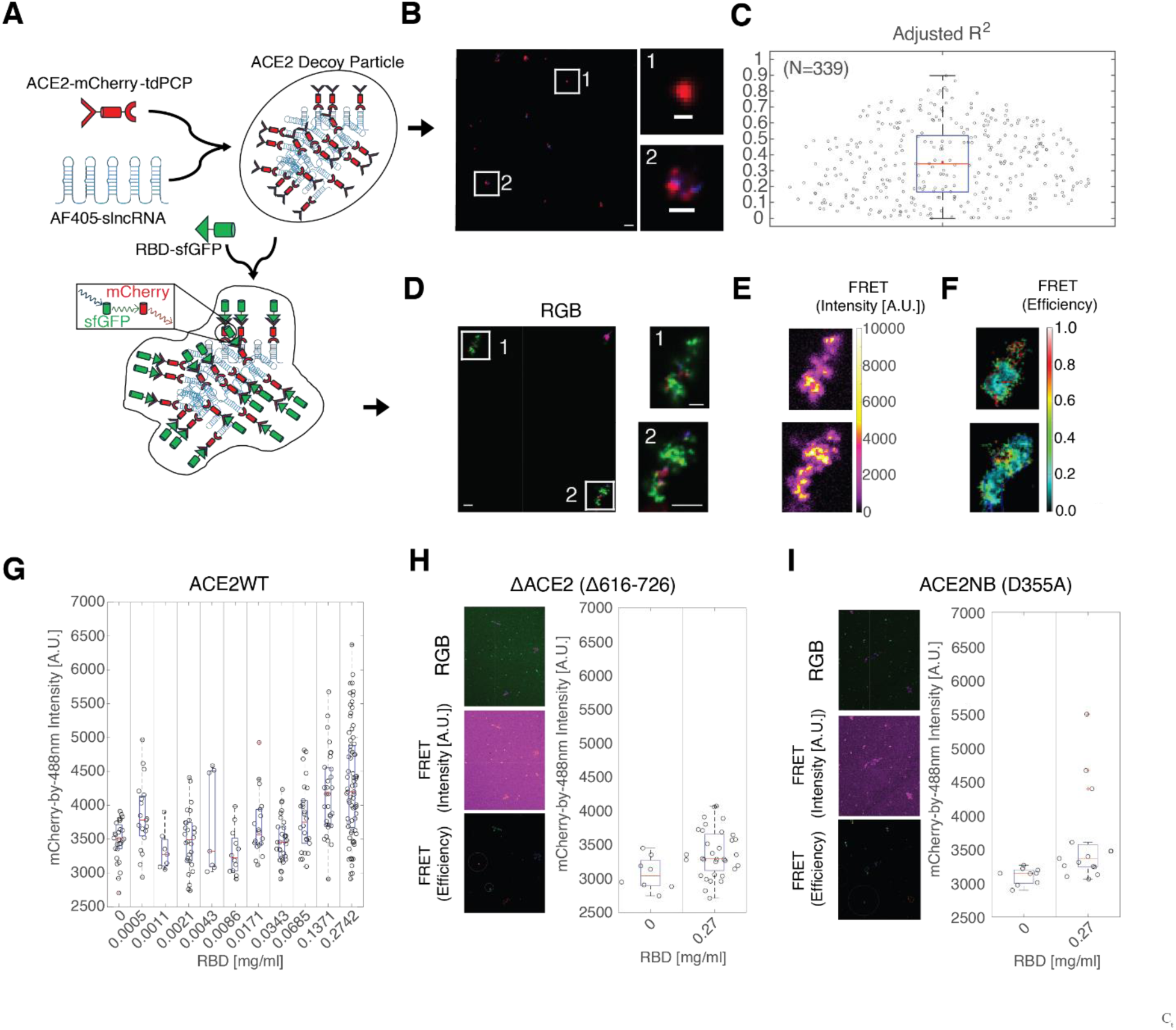
ACE2 decoy granules exhibit FRET excitation with SARS-CoV-2 RBD. **(A)** Scheme of assay: ACE2 labelled with an mCherry fluorescent protein and fused to tdPCP is incubated with a slncRNA cassette, encoding for multiple PCP-binding hairpins. 1h post incubation, an sfGFP-labelled RBD is added, facilitating FRET with the mCherry-labelled ACE2 proteins bound to the slncRNA granules. **(B)** (Left) Representative image of ACE2 granules in a typical field of view. Scalebar: 5µm. (Right) Enlarged images highlighted by white squares on the left. (Right-top) protein only condensate. (Right-bottom) An sRNP granule of protein co-localized with slncRNA. Scalebar for enlarged images 1µm and 2µm for top and bottom, respectively. **(C)** Correlation between mCherry and AF405 signals intensities of 339 mCherry-positive events. Representative events – see 1 and 2 from panel B and Supplementary Figure 2A. **(D)** (Left) Representative image of ACE granules together with RBD (green label). Scalebar: 5µm. (Right) Enlarged images highlighted by white squares on the left showing typical triple labelled structures. Scalebar: 5µm. **(E-F)** FRET intensity signal **(E)** and FRET efficiency **(F)** measured for the highlighted granule structures shown in Figure 1d. (**G)** Putative green-red FRET excitation events obtained for various RBD concentrations. **(H-I)** (Left) Images of triple-labelled granules, FRET intensity and FET efficiency, and (right) putative intensity of FRET excitation events obtained for the non-dimerizing ACE2 (Δ616-726) (h) and the non-RBD-binding missense ACE2 mutant (D355A) (I).

To test for granule functionality, we then incubated the antiSARS2 decoy particles with the SARS2 RBD fused to super-folder GFP (sfGFP) and imaged the resultant granule structure using three separate color channels (RGB) on Zeiss LSM700 confocal microscope (Figure 1D and Supplementary Figure 2B). A close examination of the structures shows (see structures 1 and 2 to the right) dominant bright green patches with very little red and blue intermixed. In particular, the small red patches did not co-localize with the green patches but rather were found to be immediately adjacent. Due to the observed discontinued strong brightness in the green patches, we hypothesized that we may be observing patches of Forster resonance energy transfer (FRET) between the RBD-sfGFP and ACE2-mCherry-tdPCP. To check for green-red FRET, we rearranged our filters to capture red channel emissions while exciting in green (a built-in mode within the LSM700 confocal microscope). The resultant images (Figure 1E) exhibit a strong FRET intensity signal all across the structures, indicating strong binding and close proximity of the RBD to the ACE2. We then analyzed the FRET efficiency across our structrures and found a value of 0.2 to 0.3, corresponding to an average molecular separation of 6-8 nm between the fluorescence moieties of the RBD and ACE2 molecules (Figure 1F). We next studied the FRET signal that resulted from biocondensates incubated with an increasing concentration of RBD-sfGFP (Figure 1G). We found a small amount of FRET signal for low concentrations (below 0.0685 mg_RBD_/ml). However, a strong FRET signal emerged (i.e. above sfGFP-only basal FRET intensity) when the RBD:ACE2 molecular ratio was 3:2 (0.0685 mg_RBD_/ml). Interestingly, this trimer:dimer ratio is the natural state of binding of SARS-CoV-2 spike protein and ACE2(26). Since in this case the RBD-sfGFP does not contain the heptad repeat domains required for spike trimerization, our results suggest that optimal spike-ACE2 interaction, while strongest at a ratio of three RBDs to two ACE2, is not necessarily related to spike trimerization.

To provide additional support for our findings regarding RBD-ACE2 interaction and to test for specificity, we prepared two mutant ACE2 moieties: a variant (ΔACE2) that lacks the “neck” domain responsible for dimerization(27) (Figure 1H and Supplementary Figure 2B), and a variant with a missense mutation (ACE2NB) that abolishes RBD binding(28) (Figure 1I and Supplementary Figure 2B). The two mutant ACE2 variants formed tightly-bound granules (Supplementary Figure 2C), as was observed for the dimerized ACE2, but we did not observe colocalization of RBD-sfGFP, nor any meaningful FRET signal. Consequently, the strong ACE2-spike interaction is dependent on dimerization of ACE2 and the binding of at least three RBD moieties.

### Synthetic recombinant sialoprotein candidates form sialogranules in the presence of slncRNA cassettes

Building on the success of using the granule platform to screen for antiSARS2 functionality, we seeked do the same for influenza virus (IFV). Unlike the SARS-CoV-2 virus, which binds a single, well-defined receptor (ACE2), IFV targets post-translational modifications which are present on various host proteins and receptors. In particular, human IFV targets sialic acid (SA) attached to host proteins via an α2,6 glycosidic bond(29). SA is added to proteins in mammalian cells in various O-linked glycosylation (OLG) patterns and can also be found in two out of the three major N-linked glycosylation (NLG) patterns. To screen for antiFlu functionality, we hypothesized that phase-separated biocondensates containing slncRNA molecules and RNA-binding sialylated proteins (sialogranules) can be used to detect SA-ligand interactions. For the third molecule capable of interacting with the SA modifications (mimicking the IFV hemagglutinin [HA]), we opted to use commercially-available FITC-labeled Sambucus Nigra Agglutinin lectin (SNA), which has a stronger affinity to α2,6 SA than the IFV hemagglutinin (HA) protein(30–32). Upon introducing SNA, we expected to observe either co-localization with the granules, or transition from the granule state to another phase-separated state involving all three components due to the known agglutination properties of SNA(33, 34). A FRET signal similar to the antiSARS2 system was not likely due to the known relative weak interaction strengths between glycans and their cognate lectins(35).

To design candidate antiFlu sialogranules, we first identified a small set of putative properly sialylated proteins (i.e., possess α2,6 SA). Our tdPCP sequence itself contains two occurrences of the amino acids N-S-T, meeting the requirements for the known NLG motif N-X-S/T (N=asparagine, X≠proline, S=serine, T=threonine), which is known to be often sialylated. As a result, a granule composed from the slncRNA together with a tdPCP-mCherry moiety produced in mammalian cells (Figure 2A, left) provides a candidate sialogranule, while granules with the same slncRNA and with unsialylated tdPCP-mCherry produced in bacterial cells(36) can serve as a negative control. Given the small number of theorized sialylated sites on tdPCP-mCherry, we opted to construct an additional variant, in which the tdPCP-mCherry protein is augmented by a peptide containing five N-X-S/T sites (NXST1-mCherry-tdPCP) (Figure 2A, right). We then expressed and purified the synthetic sialoprotein variants in both mammalian and bacterial cells (HEK293F and *E. coli* TOP10, respectively, see Methods) yielding a total of four slncRNA-binding variants: tdPCPb, tdPCPm, NXST1b, NXST1m.

**Figure 2.**
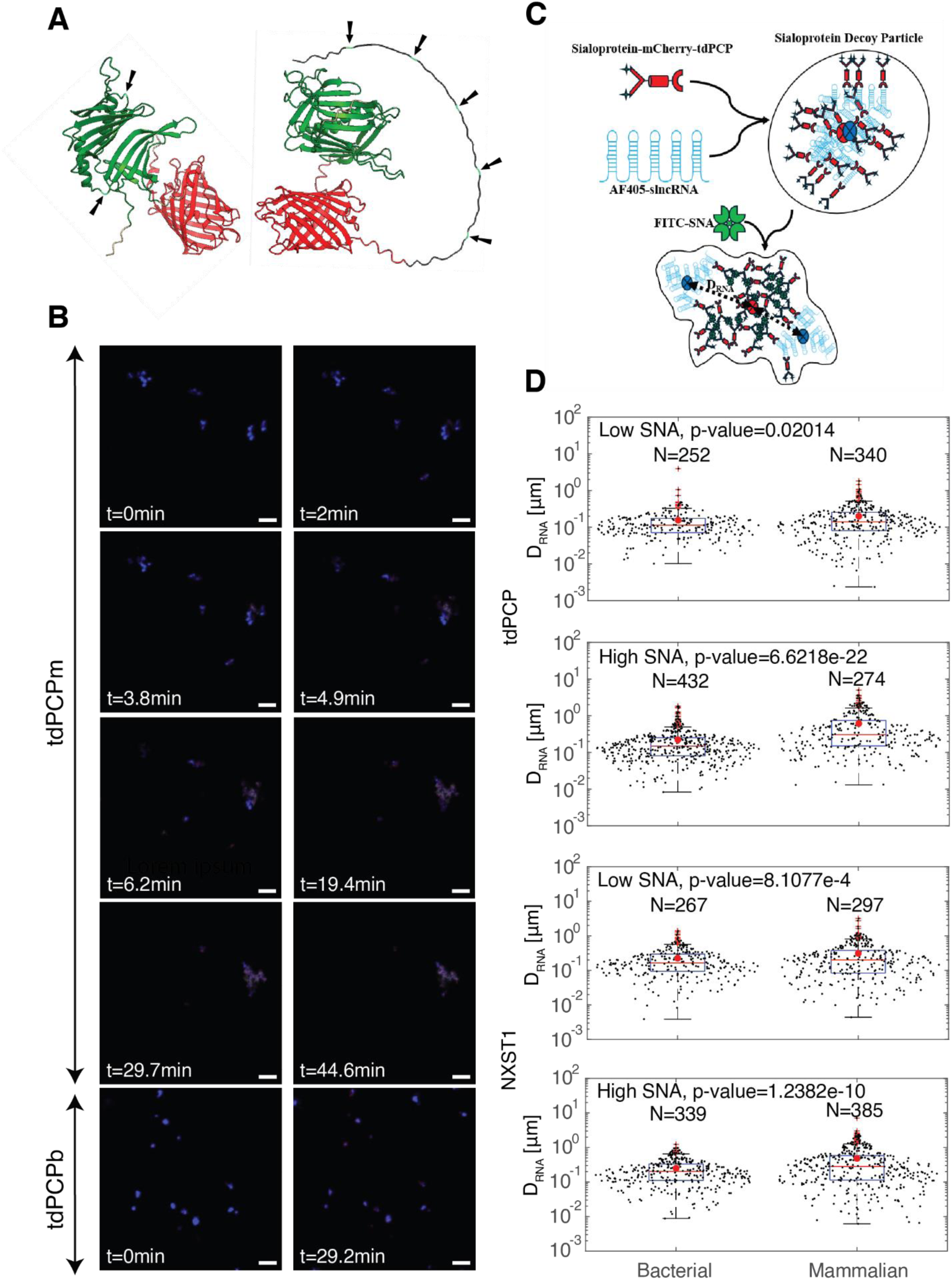
Phase separation-based agglutination assay identifies sialylated proteins. **(A)** AlphaFold structures for tdPCP (left) and NXST1 (right). Black, red, and green are NXST1, mCherry, and tdPCP, respectively. Putative sialylation sites are marked by arrows and light blow shading (asparagine residues in an N-X-S/T motif). **(B)** Accumulation of SNA signal (green) over time in a candidate sialogranules (red-blue). (Top) Candidate sialogranules composed of slncRNA (blue) and tdPCPm (red) transition to a cloudy biocondensate structure after 20min incubation with SNA. (Bottom) granules composed of slncRNA and tdPCPb (red) are not altered by SNA. Scalebar: 5µm. Images are taken from Supplementary Movies S2 and S1, respectively. **(C)** Schematic model for the transition from a sialogranules (top) to a three-component sialo-biocondensate, where the sialoprotein and SNA form a dense liquid-like phase in the center, while the slncRNA segregates to the periphery of the structure over a distance (*D_RNA_*). **(D)** Distributions comparing *D_RNA_* as defined in panel (C) for granules composed of bacterial- (left) and mammalian-(right) expressed proteins for low and high SNA concentrations. tdPCP – top two panels. NXST1 – bottom two panels.

To test for detection of sialylation, immediately after forming the granules we added the FITC-labelled SNA and observed the granules over 30min (Figure 2B). For granules containing the bacterially-produced protein variants tdPCPb, and NXST1b, no changes were observed during the 30min span of the experiment (Supplementary Movie S1). However, putative sialogranules containing tdPCPm and NXST1m exhibited an abrupt transition from a spot-like granule form to a lower-denity, cloudy biocondensate within five minutes (Figure 2B, Supplementary Movie S2). A close examination of the images and movies reveal that this biocondensate is characterized by co-localization of SNA and sialylated protein in the center of the structures, while the slncRNA is found at the periphery of the cloud (Figure 2C). As expected, no FRET signal between the FITC (SNA) and mCherry (sialoprotein) was observed (data not shown). In addition, the movies and images also reveal that the biocondensate appears to be assembled in a stepwise fashion, in a manner reminiscent of a nucleation process. Importantly, slncRNA and SNA did not colocalize without the presence of a sialoprotein (Supplementary Figure S3), and SNA-sialoprotein complexes were better detected when slncRNA cassettes were present in the reaction (Supplementary Figure S4). In the following, we refer to the composite granule-SNP as agglutinated structures.

Next, given the segregation of the slncRNA to the periphery of the agglutinated structures, we posited that we could harness the feature of the SNA-based agglutination and slncRNA exclusion process to extract a quantitative observable for differentiation between SNA-agglutinated and non-SNA-agglutinated structures. For this purpose, we define *D_RNA_* as the distance between the center of RNA clusters and putative sialoprotein clusters (Figure 2C). We then extracted *D_RNA_* from several hundred biocondensates for each granule variant, in both low and high SNA concentrations (Figure 2D). For both the tdPCP and the NXST1 proteins, a statistically-significant difference (student t-test) in *D_RNA_* distributions was observed between granules containing bacterial and mammalian components, where mean *D_RNA_* is larger for the mammalian proteins. In particular, for high SNA concentrations a much larger average *D_RNA_* is observed for both the tdPCPm (p-value=6e-22 student t-test) and NXST1m (p-value=1.2e-10 student t-test) as compared with tdPCPb and NXST1b, respectively. The increase in average *D_RNA_* is consistent with a transition from granular structure to a new type of biocondensate, characterized by a central cloud-like region that is dominated by a dense matrix of SNA and sialoprotein, while the slncRNA is segregated to the periphery of the new structure (see Figure 2C for structural model). Consequently, detection of sialic acid on granule binding protein is possible via characterization of a structural transition from the granular phase to a mixed granular-liquid phase upon introduction of SNA.

### Recombinant sialoprotein candidates form functional sialogranules in the presence of slncRNA cassettes

We next considered four natural human proteins that were either known to be sialylated or used previously in influenza research(31, 37–40): LAMP1 (Figure 3A), GYPA (Supplementary Figure S5A), fetuin (Supplementary Figure 5B), and ACE2 (Supplementary Figure S1). After fusion of the soluble domain to the tdPCP backbone, these candidate proteins contained a putative set of 21, 15, 4 and 9 α2,6 sialic acid modification sites, respectively. Since protein glycosylation is sometimes crucial for the proper folding of a protein(41), no bacterial versions of these proteins were made. Additionally, we created two additional synthetic sialoprotein candidates, comprised of tandem repeats of the NLG sequon: NXST2m, similar to the previously-described NXST1 but with different amino acids in the X position (see Supplementary Figure S5D), and tdNXT1m, a tandem repeat of NXST1 (Supplementary Figure S5C). Once fused to tdPCP, these two additional synthetic sialylated protein candidates contain 7 and 12 theoretical sialylated sites, respectively. All of these additional candidates formed granules when incubated with the slncRNA, and formed cloud-shaped biocondensate structures in the presence of SNA that were similar to the structures observed for tdPCP and NXST1 (i.e. where the RNA was found to cluster at the periphery, Figure 2), indicating that a similar phase separation mechanism is likely acting on all systems (Figure 3B, Supplementary Figure S6, and Supplementary Movie S3).

**Figure 3.**
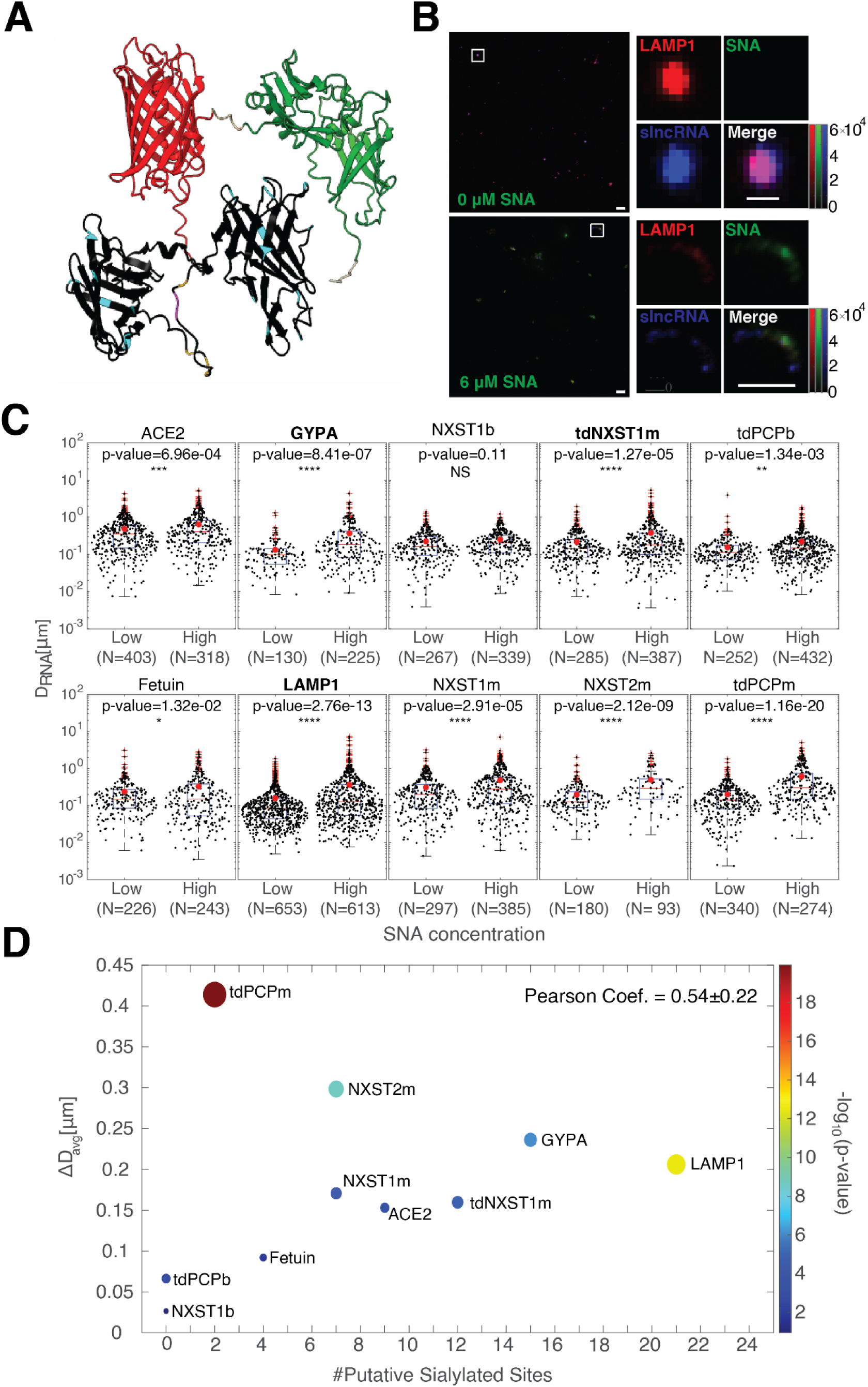
SNA-dependent formation of sialo-biocondensates. **(A)** Predicted AlphaFold structure for LAMP1-mCherry-tdPCP. Black, red, and green are LAMP1, mCherry, and tdPCP, respectively. Putative sialylation sites are marked by light-blue shading (asparagine residues in an N-X-S/T motif), yellow (serine residue), and magenta (threonine residue). **(B)** Typical field-of-view of SNA-LAMP1 granule biocondensate divided into the individual channels. (Top-left) protein (mCherry). (Top-right) SNA (AF488). (Bottom-left) slncRNA (AF405). (Bottom-right) Merged panel with all three images overlayed. (Top-set) No SNA added (scalebar: 5µm FoV, 1µm enlargement). (Bottom-set) High SNA concentration (scalebar: 5µm). **(C)** Box-plot distributions for *D_RNA_* for each candidate sialoprotein. The mean of each distribution is marked by red circle. Low and high SNA concentrations were set at below 1µg/ml (molar ratio of ∼1 SNA : 50 sialoproteins) and above 50µg/ml (molar ratio of ∼3 SNA : 2 sialoproteins), respectively. **(D)** Plot showing ΔD_avg_ as a function of putative sialylated sited. ΔD_avg_ is defined as the difference between the high and low mean *D_RNA_* values obtained for each protein in panel C. Color and size of dot correspond to the student t-test p-value computed for each panel in Figure 2C.

We then quantified the *D_RNA_* values for several hundred biocondensate structures at low and high SNA concentrations, for all of the additional candidate sialylated proteins (Figure 3C). The results show strong statistically significant differences in the *D_RNA_* distribution for LAMP1 (p-value=2.76e-13 student t-test), NXST2m (p-value=2.12e-9 student t-test), and GYPA (p-value=8.41e-7 student t-test). Three proteins exhibited a moderate statistically significant difference in *D_RNA_* distribution with a low degree of statistical significance: NXST1m (p-value=2.91e-5, student t-test), tdNXST1m (p-value=1.27e-5, student t-test) and ACE2 (p-value=6.96e-4, student t-test). Finally, fetuin was found to have a weak statistical significance (p-value=1.32e-2, student t-test) and small *D_RNA_* value, and NXST1b was found to have no statistically significant difference in *D_RNA_* distribution (p-value=0.11, student t-test) between the low and high SNA concentrations (Figure 3C). We note that fetuin exhibited structures in which the RNA was not co-localized with the protein and was rather typically found in the periphery even without the presence of SNA, potentially masking any change in *D_RNA_* distribution that could be attributed to SNA (Supplementary Figure S5B and Supplementary Figure S6, top-middle). Interestingly, tdPCPm presented the highest displacement of RNA at the highest statistical significance (with significance 17 orders of magnitude higher than its bacterial counterpart).

Finally, we plotted the difference between low and high concentrations of mean *D_RNA_* (𝞓D_RNA_) for each protein in our set as a function of number of putative sialylated sites (Figure 3D) and found them to be strongly correlated (Pearson coefficient = 0.54). Interestingly, tdPCPm and NXST2m were outliers, exhibiting a much larger 𝞓D_RNA_ response as compared with the general trend, suggesting that the number of sialylated sites or sites that are available for interaction may be influenced by other features(42). In addition, both tdPCP and NXST1 exhibited a ∼6-fold difference in 𝞓D_RNA_ between the mammalian and bacterial isoforms (tdPCP: bacterial=66.2nm, mammalian=414nm, NXST1: bacterial=26.4nm, mammalian=170.6nm), providing further evidence that the underlying response was dependent on sialylation.

### Kinetics of both SNA binding to sialogranules and slncRNA exclusion exhibit a dependence on the number of putative sialylation sites

Given the correlation observed between the number of putative sialylated sites and 𝞓D_RNA_, we wondered whether the kinetics of the phase-separation-based response to SNA concentration also correlate with the theoretical SA-content of the sialoprotein candidates. To check this, we mixed a pre-formed fixed amount of granules with a set of ten increasing SNA concentrations, and analyzed the phase-separated structures that formed. In brief (see Methods), for each of the sialoprotein candidates (see Figure 4A for a sample field of view), we first identified mCherry-positive clusters of pixels (events) that were higher than a predetermined threshold (Figure 4B). Each event was considered colocalized with slncRNA if it shared sufficient amount of pixels with at least one AF405-positive event in its periphery, colocalized with SNA if 70% of its pixels area was above a threshold for FITC (Figure 4B-top quad), or colocalized with both (triple labelled) if both criteria were met (Figure 4B-bottom quad). Each of these colocalizes structure-types would appear in some frequency in a typical image, as well as some events deemed not colocalized, for which no AF405 or FITC could be detected above their respective thresholds.

**Figure 4.**
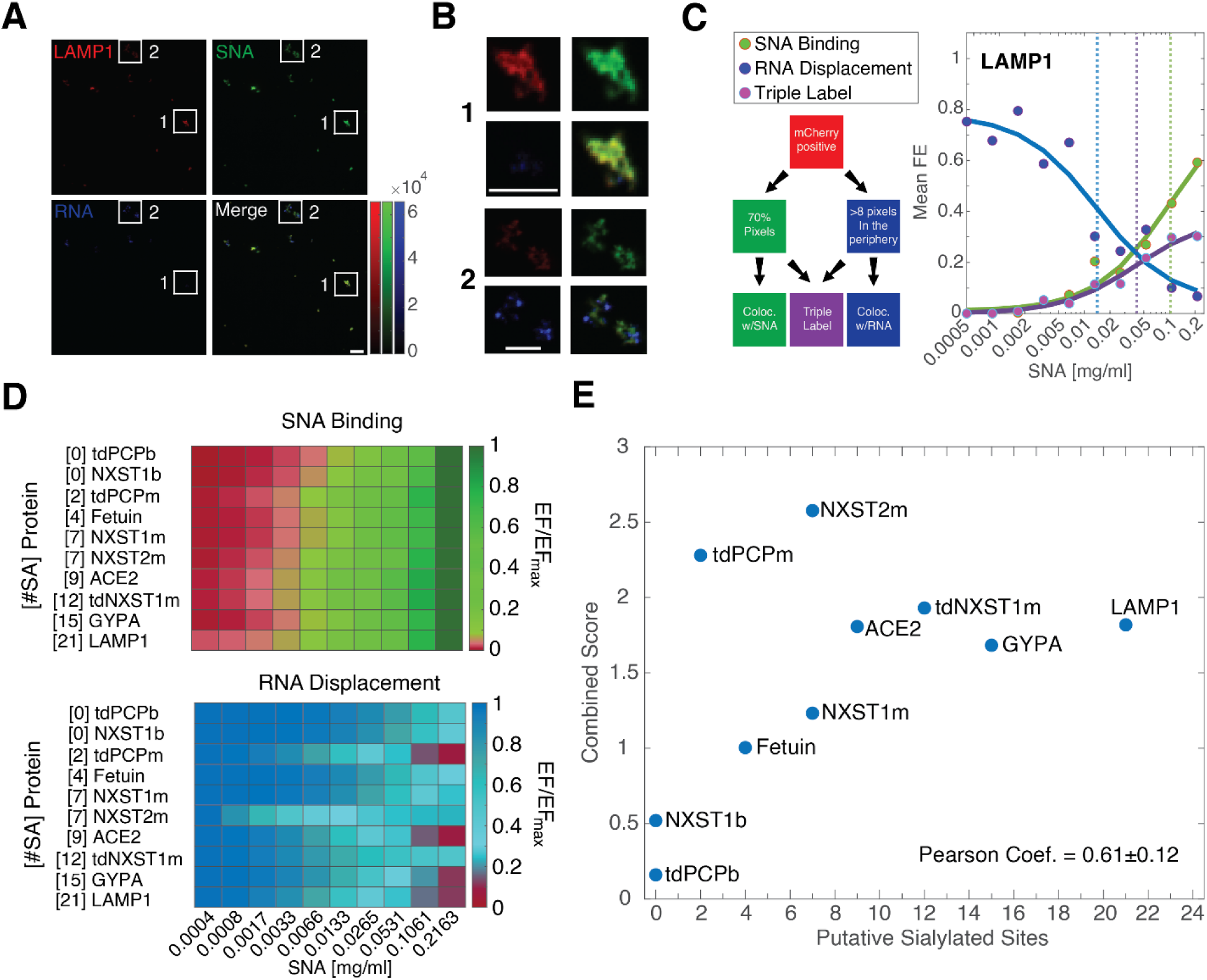
SNA-granule biocondensate formation depends on the number of sialylated sites. **(A)** Typical field-of-view of SNA-LAMP1 granule biocondensate divided into individual channels. (Top-left) protein (mCherry). (Top-right) SNA (AF488). (Bottom-left) slncRNA (AF405). (Bottom-right) Merged panel with all three images overlayed. Scalebar: 5µm. **(B)** Close-up on the two features highlighted in panel a displayed via the same three separated channels and merged image. Scalebar: 5μm. **(C)** LAMP1 dose-response curves for RNA-displacement (blue), SNA binding to protein (green), and triple colocalized biocondensate structures (purple). (Left) decision tree used to classify the different structures. **(D)** Dose responses (see Supplementary Figure S7B) displayed as a normalized heatmap for SNA binding (top) and RNA displacement (bottom). Each row was normalized by its maximum value. **(E)** Plot of the combined agglutination score for each candidate sialoprotein as a function of the number of putative sialylation site showing a strong linear dependence. Each score is the summation of three separate values: ΔD_RNA,_ ΔΔG_SNA binding_, and ΔΔG_RNA displacement_ (see methods and text for definitions).

The analysis shows (Figure 4C and Supplementary Figure S7) that for almost every putative sialylated protein, an increase in the SNA-sialoprotein co-localized events as well as SNA-sialogranule triple-labelled spots are observed for increasing SNA concentration, while simultaneously the frequency of SNA-lacking slncRNA-sialoprotein declines. The mean event frequency (EF) of each colocalization type were then fitted as increasing or decreasing Hill curves for either SNA binding or RNA displacement, respectively (see curves in Figure 4C and Supplementary Figure S7B). To compare between the different candidates, each fitted event frequency was normalized by the maximal event frequency of the respective sialoprotein (Figure 4D). For SNA binding, the results show a dependency on the number of putative sialylated sites, where a significant FITC signal could be detected at lower SNA concentrations for higher putative number of sialylated sites (Figure 4D, top). For RNA displacement, gradual decrease in AF405 signal as the SNA concentration increases. Interestingly, both tdPCPm and NXST2m display a slightly outlying behavior, where AF405 signal is displaced at relatively low SNA concentrations, similarly to what was observed in 𝞓*D_RNA_* measurements (Figure 4C, bottom). We extracted Hill constants (*K_d_*) from the fitted curves for each of the sialoprotein candidates, and normalized by the *K_d_* for tdPCPb, to estimate an effective free energy for SNA-granule biocondensate structure formation (𝞓𝞓*G*, see Methods). For both SNA binding and RNA displacement, 𝞓𝞓*G* values correlated strongly with the number of putative sialylated sites (Supplementary Figure S8), with similar Pearson correlation values to what was measured for the 𝞓*D_RNA_* measurement (Figure 3D).

Finally, as a proposed score of sialylation, we considered 𝞓*D_RNA_* values and the 𝞓𝞓G values for both SNA binding and slncRNA displacement as three independent measurements. We ranked the different candidate sialoproteins (Figure 4E) by normalizing the result for each measurement by its respective maximal value, and summing the normalized values (see Methods). We observe a strong Pearson correlation (0.61) between the proposed sialylation score and the number of putative sialylated sites, with only tdPCPm and NXST2m exhibiting outlier behavior. Altogether, the SNA dose-response analysis provides further support that SNA-mediated phase transitions can differentiate between sialylated and unsialylated proteins, and in addition also suggests that a reliable estimate of the number of sialylated sites on a target protein may be obtained from such a phase-separation-based assay.

### Decoy granules inhibit influenza infection using a pinball-like disrupted diffusion mechanism

To test whether the phase-separation-based responses of the granules to their respective virus-capsid-mimicking third molecular components reflect actual antiviral activity, we opted to employ a viral entry assay for IFV and SARS-CoV-2 (Figure 5A). Given the complexity of the potential outcomes, we first simulated a simplified infection process that we term “disrupted diffusion” (Figure 5B, top left schema). We hypothesized that a diffusion process(43) controls the underlying dynamics of the virion search for its cellular target. Adding the (larger) decoy granules to the mix effectively introduces quasi-static obstacles positioned at random locations in space. Therefore, in this simplified scheme, the decoy granules serve as large static obstacles, where the smaller dynamic virions bind transiently and then bounce to the next decoy target in a stochastic pinball-like manner, leading to a disrupted diffusion process (Figure 5B, bottom right schema). The results show (Figure 5B) that the disrupted diffusion leads to an increase in the average diffusion time per virion (Supplementary Figure S9A and S9B), which in turn translates to a reduction of the virion flux on the cell layer, resulting in inhibition of infection. This model is consistent with the current understanding of IFV infection(31, 44, 45), where upon engaging with large natural soluble sialoproteins (e.g. mucins) the IFV neuraminidase protein cleaves sialic acid, and thus consistently disrupts any hemagglutinin (HA)-SA binding events and prolongs IFV diffusion time(46–48) and affects IFV trajectory(31, 45). The simulation further shows that as the number of sialylated decoy particles or “obstacles” per volume increases, the flux of the virions on the simulated cell layer decreases (Figure 5B and Supplementary Figure S9B), as expected. In addition, increasing the number of sialic acid modifications, which is implemented in the simulation as increased duration of interaction between virions and obstacles, leads to a further decrease of the virion flux (5b, bottom and Supplementary Figure S9B). Conversely, a simulation of a suspension containing only sialoproteins that do not phase separate, implemented as 0 obstacles (Figure 5B, top), does not meaningfully inhibit virion flux on the cell layer.

**Figure 5.**
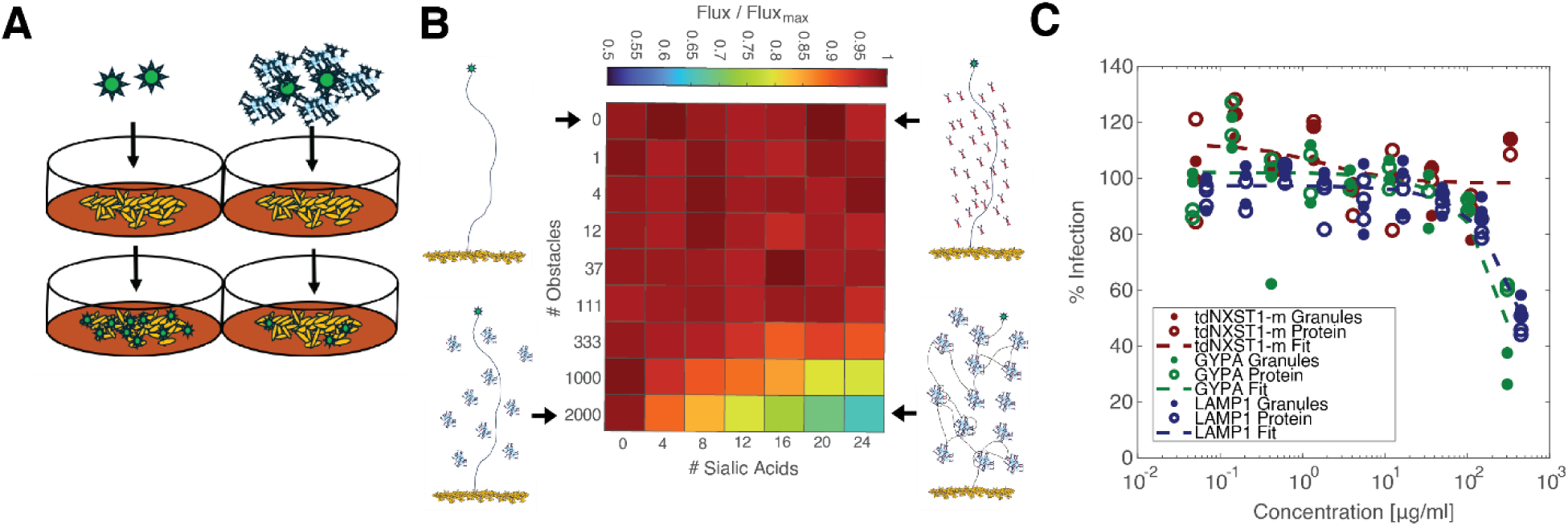
A disrupted diffusion model accounts for inhibition of IFV infection by LAMP1 or GYPA decoy particles in A549 cells. **(A)** schematic of viral entry assay. Influenza virions (green objects) infected A549 cells (yellow objects) in the absence (left) or presence (right) of sialogranules. **(B)** Normalized heatmap of virus infection flux (see Methods). Increasing the number of diffusion obstacles (y-axis) or the strength of their interactions (x-axis) results in decreased viral flux. Accompanying schemas: (top left) no obstacles. (Top right) Soluble sialoproteins. (Bottom left) Obstacles without sialic acid. (Bottom right) Obstacles with high interaction strength. **(C)** Entry assay results for three sialoprotein candidates in various sialogranules concentrations. GYPA (green) and LAMP1 (blue) were able to inhibit IFV infection, while the synthetic tdNXST1m (red) was not. Trendlines are fitted using both sialogranules and sialoproteins-only data. Results are normalized for the mean infected untreated cells in each concentration row.

We experimentally tested several antiFlu sialoprotein decoy candidates using an ex vivo influenza infection model(49). in presence and absence of the slncRNA (Figure 5C). The results show that only LAMP1 and GYPA triggered a reduction of infection consistent with the pinball model’s predictions, while NXST1m did not exhibit antiviral activity, despite having verifiable sialic acid modifications. In addition, both the LAMP1 and GYPA sialogranules (blue and green closed circles, respectively) exhibited a similar inhibition of infection to the protein-only samples, culminating in a maximum of ∼70% and ∼45% reduction at the highest granule concentrations. Since tdNXST1m exhibited smaller biocondensates in both granule and protein-only form as compared with LAMP1 and GYPA, these observations suggest that biocondensate structure and size, and not only the number of sialic acid modifications on the protein, may be important factors in viral inhibition.

### Inhibition of SARS-CoV-2 infection by granules suggests a more complex infection mechanism

We next tested the efficacy of ACE2 decoy granules against both the Delta and Omicron BA.1 SARS-CoV-2 variants via a similar viral entry assay using Vero E6 cells (see Methods). An initial scan of the infected cells revealed that antiSARS2 decoy granules demonstrated complete inhibition of infection at the highest granule concentrations (Figure 6A). A closer observation of the anti-viral response (Figure 6B) further revealed enhancement of infection at lower protein concentration, which transitioned to complete inhibition of infection at ACE2 concentrations > 3 μg/ml. In particular, the protein-only samples showed stronger enhancement (empty circles) as compared with the granules (filled circles) and transitioned to inhibition at higher protein concentration thresholds. This difference can be quantified by computing the IC_50_ value for both the granules and protein-only cases (Supplementary Figure S10, green vs blue lines). For the Delta variant, a 7-fold reduction (∼87%) in IC_50_ was recorded (3.15µg/ml and 22.9µg/ml, for the decoy granules and ACE2-only, respectively) while a ∼33% reduction was recorded for the Omicron BA.1 variant (IC_50_ = 3.1µg/ml and 4.6µg/ml for decoy granules and ACE2-only, respectively). Finally, the slncRNA elicited no antiviral response at any concentration for either variant, as expected, and weak-decoy like response consistent with the pinball model at the highest concentrations (Supplementary Figure S10, red).

**Figure 6.**
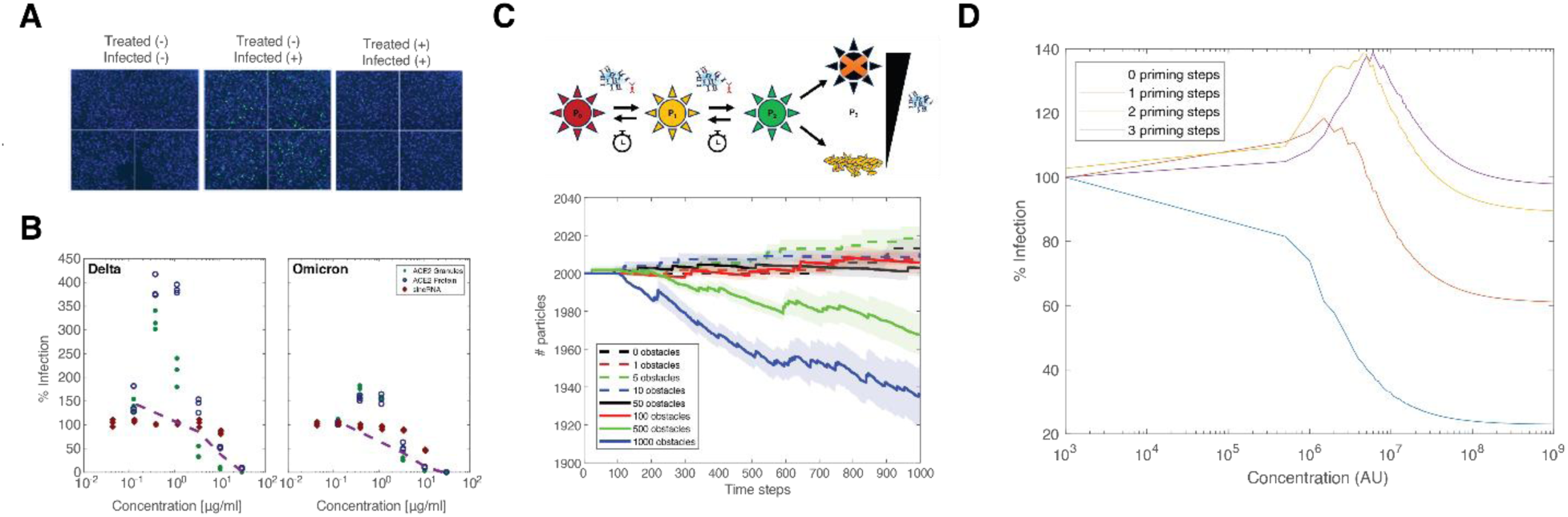
Interaction with ACE2 decoy particles enhances viral infection in low concentration and inhibits it at high concentrations. **(A)** Representative FoVs of infection by the Delta variant from the highest tested ACE2 decoy particle concentration. Blue signal – DAPI (cell). Green signal – AF488 (anti-spike antibody). **(B)** Entry assay results for either the Delta (left) or Omicron (right) variants, in various ACE2 decoy particles concentrations. Below 3µg/ml, ACE2 decoy particles demonstrate enhancement of infection, while above viral infection is fully inhibited. Trendlines (magenta) are fitted using both particles and ACE2-only data. **(C)** Priming model schema (top) and results (bottom). Prior to cell infection, a SARS-CoV-2 virion encounters either an ACE2 decoy particle or soluble ACE2, affecting one of its RBDs. A third priming event results in infection (if done in the cell layer) or virus elimination (if done by a decoy particle). Without decoys or free proteins, overtime the priming is reversed. At low obstacles numbers (<100, a virus:obstacle ratio of 20:1), viral titer increases, while higher number of obstacles reduce the viral titer. **(D)** The model described in equations 10-15 was solved via MATLAB for multiple primed state with the following values for the constants: 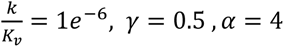.

The experimentally-observed enhancement of infection at lower protein titer was not observed in our initial simulations using the simple pinball model. In an attempt to provide a simple biological mechanism, we added a “priming” step to the disrupted diffusion simulation. We define priming as an effect that occurs prior to virus activation. In this case, at least one ACE2-spike binding event must occur at either an obstacle or the cell wall, for the virus to become active, or primed, for infection at a subsequent interaction with the cell wall. Qualitatively, in the case of a low concentration of obstacles, a virion may encounter one or two obstacles before reaching the cell surface. This implies that while the flux of virions on the cell wall will decrease somewhat, as expected due to the presence of the obstacles, the actual flux of primed virions may increase over some low range of obstacle concentration, leading to enhancement of infection (Figure 6C). To simulate priming, we set up kinetic equations where the virus transitions from unprimed to as many as 3 primed states in a stepwise fashion (see model section in Methods). The simulations for various number of priming steps (Figure 6D) show that at two priming steps and above, enhancement of infection occurs in a fashion that is similar to what was observed in the SARS-CoV-2 infection experiments. Interestingly, the model indicates that a higher of number of priming steps leads to a stronger enhancement effect, suggesting that the difference between the antiviral response recorded for Omircon and Delta variants may stem from the latter requiring one more priming step than the former. Consequently, the inhibition patterns generated by the various decoy antiviral granules seem to reflect different underlying molecular and kinetic mechanisms associated with infection.

## Discussion

The COVID-19 pandemic revealed that our antiviral therapeutic toolbox is woefully lacking broad-spectrum solutions that can be deployed rapidly to combat any emergent infectious disease with pandemic or endemic potential. To address this lack, we propose synthetic RNA-protein (sRNP) granules that can function as a broad-spectrum decoy therapeutic, which prevents infection by binding invading virions, inhibiting them from infecting cells. Our synthetic RNA-protein (sRNP) granule platform carries out two critical research functions. First, granule formation and phase separation can be utilized to quantify a wide range of biomolecular interactions, either via colocalization or via structural changes to the granules. Second, the interaction metrics can be utilized to optimize sRNP granule composition, and to screen for functional protein components. The most promising formulations can subsequently be tested for antiviral decoy activity via traditional viral entry assay, that is both expensive and often requires a higher biosafety-level facility. Our results show that the granule platform enabled detection of both ACE2-SARS2-RBD interaction, and of various sialylated proteins via interaction with SNA lectin. In particular, the strong ACE2-RBD interaction led to the formation of gel-like particles consisting of three components, where a FRET signal between the ACE2 and the RBD proteins suggested an average displacement of ∼6-8 nm between proteins. By contrast, the weaker lectin-sialic acid interaction led to the “melting” of the dense granules into a liquid-like phase in which both protein components were localized, while the RNA component segregated to the periphery of the new structure. The viral entry assays confirmed the antiviral potential of ACE2, GYPA, and LAMP1 granules. The proposed sRNP platform opens the door for the development of a true broad-spectrum decoy antiviral, comprising of a slncRNA cassette and potentially any number of slncRNA-binding soluble receptors. Finally, by incorporating different RNA binding protein moieties and their respective binding sites, this platform can in theory be molecularly programmed to inhibit single or different combinations of virus species.

A deeper examination of our results allows us to make additional observations. The structural transition phenomenon and its correlation to the number of sialic acid modifications (Figure 3D and 4E) on the target protein is consistent with polyphasic linkage theory(18), which provides a biophysical mechanism for the regulation of biomolecular condensates by ligands. As a result, this regulatory effect should be generalizable to other post-translational modifications, and the structural melting of granules can form the basis of a new type of “binding” assay. Such an assay can be employed with relative ease, and could serve as an alternative to existing ELISA-based or mass spectrometry-based diagnostic approaches. In particular, our ability to differentiate between two genetically-identical protein products based solely on the presence of post-translational sialylations due to the cell-type in which they were expressed provides a simple proof-of-concept as to the validity and utility of this assay (Figure 2D).

Moreover, it is also worth noting that consistent with the fact that the RBD-granule binding assay and the lectin-granule binding assay generated different structural disruptions to the granule platform, the antiFlu and antiSARS2 granules also exhibited different antiviral responses. While the antiFlu GYPA and LAMP1 granules inhibited IFV infection only at high protein concentrations, consistent with a simple disrupted-diffusion or “pinball” model (Figure 5C), the antiSARS ACE2 granules showed a more complex antiviral behavior (Figure 6B), suggesting a more nuanced receptor-RBD interaction. While enhancement of infection by SARS2 variants at low ACE2 concentrations is a controversial subject and has only been observed in very few reported studies(50–52), the results shown here provide an impetus for further research into this phenomenon to confirm whether or not it is a real feature of the SARS2 infection process or an artifact of the Vero assay used here and elsewhere(50, 51). In particular, the proposed mechanism of virus priming (Figure 6) is consistent with several structural analysis studies that were carried out on ACE2-spike complexes(53–60). Therefore, while it is entirely possible that the simplified infection models devised here may only be relevant to the particular cell-types and entry assays used in our experiments, they may shed light on complex kinetic phenomena that play a crucial role in both SARS-CoV-2 and IFV infection. Consequently, our platform constitutes not only a new potential broad-spectrum therapeutic, but also provides an alternative assay for the discovery and study of viral infection processes, which could ultimately yield new antiviral therapeutic strategies.

## Materials and Methods

### Plasmid construction

Protein genes were ordered as either Gene Fragments (Twist Bioscience) or gBlocks (Integrated DNA Technologies, IDT). For the natural proteins LAMP1, GYPA, Fetuin/AHSG and ACE2, the sequence that was cloned was extracted from UniProt and contained the protein’s native signal peptide and the extracellular domain, namely amino acids 1-382, 1-91, 1-367, and 1-740, respectively. For the NXST peptide variations, the signal peptide of ACE2 was used. The NXST peptide is comprised of five alternating repeats of the known sequon for N-linked glycosylation N-X-(S/T) with six amino acids separating between each sequon. For the mammalian tdPCP protein, the signal peptide of ACE2 was used directly upstream to the protein sequence. pcDNA3-SARS-CoV-2-S-RBD-sfGFP was a gift from Erik Procko (Addgene plasmid #141184), and a his-tag was added to the C-terminus of the RBD-sfGFP open reading frame (ORF). 𝞓ACE2 was created by removing amino acids 616-726 from the ACE2 ORF. ACE2NB was created by replacing the codon for D355 with a codon for alanine. All sequences can be found in Supplementary Table 1.

All but two proteins were expressed in HEK293F cells. Of the proteins produced in mammalian cells, all but RBD-sfGFP were made from genes cloned into a pTwist-CMV-BetaGlobin vector downstream of the β-globin intron. An affinity tag comprised of six histidines (his-tag) for purification purposes was positioned immediately upstream to the stop codon of all coding sequences. Bacterial tdPCP (tdPCPb) was cloned into A133 plasmid under N-butyryl-L-Homoserine lactone (C4HSL) induction and expressed in TOP10 *E. coli* cells. NXST1b was cloned into A133 plasmid using the coding region from the pTwist-CMV-BetaGlobin vector, and expressed in the same manner as tdPCPb. Plasmids containing the synthetic long non-coding RNA (slncRNA) cassettes were created as previously described(11).

### In vitro transcription and purification of RNA cassettes

RNA transcription of synthetic RNA cassettes was done as previously described(11). Briefly, a vector containing the slncRNA DNA sequence, flanked by two EcoRI restriction sites, was digested with EcoRI-HF (NEB, cat. R3101L) per the manufacturer’s instructions to form a linear fragment. The enzyme was then heat-inactivated by incubating the restriction reaction at 65 °C for 20 min. For fluorescently labeled RNA, 1µg of the restriction product was used as template for in vitro transcription using HighYield T7 Alexafluor-405 (AF405) RNA labeling kit (Jena Bioscience, cat. RNT-101-AF405), according to the manufacturer’s instructions. Non-fluorescent RNA was transcribed using the HiScribe™ T7 High Yield RNA Synthesis Kit (NEB, cat. E2040S). Following in vitro transcription by either kit, the reaction was diluted to 90 µl and was supplemented with 10 µl DNAse I buffer and 2µl DNase I enzyme (NEB, cat. M0303S) and incubated for 15 min at 37 °C to degrade the DNA template. RNA products were purified using Monarch RNA Cleanup Kit (NEB, cat T2040S) and stored in −80°C.

### Cell maintenance

A cryotube with 1ml (10%v/v DMSO (Sigma-Aldrich, cat. D2650)) of 10^7^ HEK293F cells/ml (Thermo Fisher, cat. R79007) was thawed as follows: the cryotube’s content was transferred into 4ml fresh Freestyle media (Thermo Fisher, cat. 12338018), prewarmed to 37°C, and then immediately centrifuged at 500g for 5min. Supernatant was removed, and cell pellet was resuspended in 7ml of fresh Freestyle media and transferred to 23ml of prewarmed Freestyle media in a 125ml flat-bottom flask (TriForest, cat. TF FPC0125S). Cells were then moved to an incubator and grown under 37°C, 8% CO_2_ and humid conditions (an in-hood basin with 500ml autoclaved DI water, supplemented with 1%v/v Aquaguard-1 solution (Sartorius, cat. 01-867-1B) before use). The cells were placed on an in-incubator orbital shaker which rotated at 135rpm. These conditions were in accordance with the cells’ manual.

The cell concentration exceeded 1.2x10^6^ cells/ml 3-4 days after thawing, at which point cells were passed by diluting them 1:5 in a final volume of 30ml fresh growth media. Cells were routinely passed every 3-4 days, as cell concentration was sufficient. Cells were checked for mycoplasma in accordance with manufacturer protocol (Vazyme, cat. D101-02).

### Cell transfection

24h before transfection, HEK239F cells were seeded at 0.6-0.7x10^6^ cells/ml in 30ml fresh Freestyle media and allowed to grow overnight. On the day of transfection, cells were diluted to 1x10^6^ cells/ml.

Per flask containing 30ml of culture, up to 40µg of plasmid DNA was diluted in OptiMEM buffer (Thermo Fisher, cat. 31985070) to a final volume of 600µl. 120µl of 0.5mg/ml branched PEI solution (Sigma-Aldrich, cat. 408727) or 60µl of 1mg/ml linear PEI (Polysciences, cat. 23966-1) was diluted in OptiMEM buffer to a final volume of 600µl. The PEI-OptiMEM solution was added to the DNA-OptiMEM solution (final mixed volume of 1.2ml) and incubated at room temperature for 15min. Each flask was then transfected with the appropriate plasmid within 5 minutes. Transfected cells were maintained for 5-7 days, during which time the growth media changed color to red due to the mCherry-label in the recombinant proteins.

### Protein extraction from HEK293F cells

1ml of nickel (Ni)-coated beads (Promega, cat. V8821) were transferred into 50ml conical tubes and were allowed to settle. 400µl of supernatant was removed and beads were washed in 2.5ml of equilibration buffer (50mM NaH_2_PO_4_, 300mM NaCl, 5mM imidazole, pH 8). After the beads resettled, 2ml of the supernatant was removed.

Transfected cells were centrifuged 5-7 days post transfection at 5000rpm for 20min at 4°C. Up to 40 ml supernatant was transferred to a 50ml conical tube with settled beads while cell pellet was discarded. Bead-protein mix was incubated at room temperature for 1h in an end-over shaker. After incubation, the solution was transferred to a gravity column (Bio-rad, cat. 731150) and ∼100µl of the flowthrough was collected. Next, beads were washed three times using 5ml of wash solution (50mM NaH_2_PO_4_, 300mM NaCl, 20mM imidazole, pH 8), and ∼100µl was collected from the first wash. Finally, beads were resuspended inside the column in 2ml elution buffer (50mM NaH_2_PO_4_, 300mM NaCl, 500mM imidazole, pH 8) and left to incubate for 15min, after which the entire elution volume was collected in 2-3 fractions.

### Protein extraction from TOP10 *E. coli* cells

TOP10 *E. coli* cells (Invitrogen, cat. C404010) were grown in 5ml Luria broth (LB), supplemented with 100µg/ml ampicillin (Sigma-Aldrich, cat. A9518), at 37°C and 250rpm overnight. The following day, the starter content was transferred to 500ml TB (24g yeast extract, 20g tryptone, 4ml glycerol, in 900ml DI water), supplemented with 100µg/ml ampicillin and 100µg/ml C4HSL (Cayman Chemical, cat. 10007898), and grown overnight at 37°C and 250rpm. The next day, cells were allowed to grow until reaching an optical density of >0.45, before being centrifuged at 6000rpm for 10min. Supernatant was removed, and cell pellet was resuspended in 30ml of resuspension buffer (50mM Tris base pH=8, 100mM NaCl, 0.02% sodium azide, in 1L DI water, titrated to pH=7), before being passed four times through an EmulsiFlex-C3 homogenizer (Avestin Inc.). Finally, cellular debris was centrifuged at 13000rpm for 30min at 4°C and supernatant was transferred into a new 50ml conical tube. Purification of proteins were executed as described for proteins produced in HEK-293F cells, after adding up to 40ml clarified cell lysate to settled Ni-coated beads.

### Verification of extracted proteins using SDS-PAGE

15µl of each fraction (flowthrough, wash, elution) per protein was taken to a sodium dodecyl sulfate polyacrylamide gel electrophoresis (denaturing SDS-PAGE), using a 10% acrylamide concentration in the resolving gel. 0.1mg/ml of purified BSA (NEB, cat. B9000) was used as positive control. Samples were boiled at 95°C for 10min before loading onto the gel. Gel running conditions were 150V for 70min. Afterwards, the gel was stained using Instant Coomassie Blue (Abcam, cat. Ab119211) for 15min.

### Buffer exchange of extracted proteins

The high imidazole content from the elution buffer was diluted ∼1:10^4^ by using an Amicon filtration tube (Millipore, cat. UFC200234) according to the manufacturer’s protocol. In brief, PBSx10 was added to the eluted protein samples as 1/9 volume equivalent its post-extraction volume. Then, PBSx1 was added to dilute the imidazole concentration to 100mM, the maximal concentration recommended by the manufacturer’s protocol. The diluted protein solution was loaded onto the Amicon tube and centrifuged for 15min at 5000rpm several times, until reaching 400-500µl retentate volume. In between each centrifuge, protein precipitate was resuspended by carefully pipetting inside the column. Once arriving at the appropriate volume range, 500µl of PBSx1 was added to the protein solution inside the column, diluting the imidazole ∼2-fold. This process was repeated eleven times.

### Formation of RNA-protein granules

For microscopy assays, 250fmol slncRNA PP7x8 was mixed with any of the proteins at a molar ratio of 100:1 protein-to-RNA molecules. For agglutination assays and viral entry assays, proteins and slncRNA were incubated in 7µl of granule solution (750mM NaCl, 1mM MgCl_2_, 10% PEG 4000) and completed with water to a final volume of 21ul per reaction. For FRET assay, ACE2 and slncRNA were incubated in 15µl granule buffer (1M NaCl, 1.33mM MgCl_2_, 13.3% PEG 4000, 100mM KH_2_PO_4_, pH=7.44) and completed with water to a final volume of 20µl per reaction The mixture was incubated at room temperature for 1h before being taken for imaging. For viral entry assays, the same protocol was maintained with a protein:slncRNA ratio was 10:1, and the slncRNA was PP7x14-MSx15 cassette.

### Confocal microscopy imaging

For all imaging, we used the LSM-700 (Carl Zeiss) confocal microscope, equipped with a BIG (GaAsP detector) unit, using a Plan-Apochromat 63x/1.40 Oil objective. Images were captured using Zen imaging software (version 8.0, 2012) with a FRET module. A pinhole of 1.5µm was set for each laser. Emission windows were 578-800, 300-583, and 300-483 for the 555nm (mCherry), 488nm (FITC), and 405nm (AF405) channels, respectively, for all assays other than the FRET assay. Other parameters (e.g. gain, intensity, averaging time, etc.) varied between assays, based on the acquisition and assay needs, and are stated below for each microscopy assay. Other than the time lapse experiments, for each assay we used standard microscopy slides with 1.5H coverslips, and each sample was imaged using 5µl.

### ACE2-RBD colocalization and FRET analysis

5µl of ACE2 granules was mixed with 1µl of RBD-sfGFP (either from the stock concentration of 1.782mg/ml or nine other 2-fold serial dilutions), making the highest RBD:ACE2 ratio 5.6:1. Samples were then imaged. Laser intensities were 0.6% for all dyes. Gain values were 650, 650, 650, and 850 for mCherry (555nm, 560-800 excitation window), FITC (488nm, 300-550 excitation window), FRET (488nm, 560-800 excitation window), and AF405 (405nm, 300-483 excitation window), respectively. To optimize time, an averaging value of 4 and a depth of 16bit were set. RBD-only (sfGFP) and granules-only (mCherry and AF405) controls were used to determine donor and acceptor parameters, respectively.

### Confocal time-lapse assay

16µl of granules per well were added to an Ibidi µ-slide 18-well glass bottom slide chambers (cat. 81817). After 5-10min, a field of view containing granules was located, and 1.9µl of SNA (2mg/ml stock concentration) was added, resulting in a molar ratio of ∼1 RNA : 100 sialoprotein : 560 SNA. Samples were imaged immediately after the addition of SNA, and continuously for 30-45min, at 8.61 seconds-per-frame. Laser intensities were 5%, 1%, and 3.5% for mCherry, FITC, AF405, respectively. Gain values were 700, 550, 794 for mCherry (555nm), FITC (488nm), and AF405 (405nm), respectively. To optimize time, an averaging value of 2 and a depth of 16bit were set.

### Confocal agglutination assay of RNA-protein granules with SNA lectin

2mg/ml stock concentration of FITC-labelled SNA Lectin (Vector Labs, cat. FL-1301-2) was serially diluted 2-fold nine times in biological water. 1.9µl of the appropriate SNA lectin dilution was added to 16µl of granules and incubated for 10min at room temperature. Samples were then imaged. Proteins were imaged with or without RNA. Laser intensities were 2% for all dyes. Gain values were 650, 550, 750 for mCherry (555nm), FITC (488nm), and AF405 (405nm), respectively. To optimize time, an averaging value of 4 and a depth of 16bit were set. Repeats were taken on separate days, each time acquiring between three to seven fields of view per sample. All fields of view were analyzed using MATLAB code written for the analysis (see code availability statement).

### Morphological analysis of agglutination assay

mCherry-positive events were defined as a cluster of 8 or more pixels that have an mCherry channel intensity value higher than a threshold of 4000a.u. for ACE2, Fetuin, GYPA, LAMP1, and any version of the NXST peptide, 5000a.u. for LAMP1, and 3000a.u. for both tdPCP versions. FITC-positive pixels were ones with an intensity value greater than 5000a.u., and AF405-positive pixels were ones with an intensity value of 3000 a.u.

The center of gravity (CoG) of clusters of AF405-labelled pixels that occupied the same space as mCherry-positive events were calculated. Then, the distance *D_RNA_* between AF405 CoGs and mCherry CoGs for the mCherry-labelled cluster of pixels was calculated. The calculated distances in the three lowest and three highest SNA concentrations were pooled (𝞓D_RNA_), and the means distances distribution were analyzed for statistical significance using student t-test.

### SNA lectin dose-response analysis

The thresholds used for this analysis were the same as the ones used for the confocal agglutination assay analysis. mCherry-positive events were assigned to one of four colocalization groups: (1) colocalization of mCherry, FITC and AF405 signals, (2) colocalization of mCherry and AF405 but not FITC, (3) colocalization of mCherry and FITC but not AF405, and (4) no colocalization. Assignment into the four groups was done by calculating how many of the mCherry-positive pixels were co-labelled with either FITC- or AF405-positive clusters, both, or neither. For FITC colocalization, 70% of an event’s pixels had to be FITC-positive. For AF405-positive events, mCherry-labelled events had to share at least 8 pixels with an AF405-positive events, as RNA was clustered in the periphery of events as more SNA was added. The number of events in each of the four bins was divided by the total number of events per lectin concentration per protein, to yield the mean event frequency (EF) per bin per SNA concentration for each sialoprotein. The error of the mean probability was calculated according to multinomial distribution approximation as follows (equation #1):

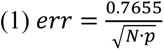

Where 𝑒𝑟𝑟 is the error, 𝑁 is the total number of observed events, and 𝑝 is the calculated probability per bin per lectin concentration per sialoprotein.

Analysis of the yielded dose response curve allowed for the extraction of dissociation constant (*K_d_*) values for either SNA binding or RNA exclusion. We used Matlab fit function for the following models (equations #2 and #3):

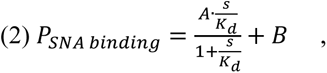

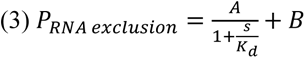

Where 𝐴 is the amplitude of response [0,1], 𝐵 is the intercept for the dependent variable 𝑃, 𝐾_𝑑_ is the dissociation constant, 𝑠 is the independent variable (SNA concentration), and 𝑃 is the probability. SNA concentration was used as mg/ml, and therefore the 𝐾_𝑑_ units are the same. Lower and upper constraint were (0,0,0) and (1,1,inf), respectively, per (𝐴, 𝐵, 𝐾_𝑑_). The fit guess for all proteins was 𝐴=maximal probability value per bin per protein; 𝐵= 0 or 𝐴 for equations #2 and #3, respectively; 𝐾_𝑑_=0.5. The fitting of the triple-labelled group was done similar to SNA binding.

Finally, calculating Gibbs free energy was performed as follows (equation #4):

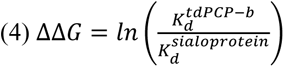

Where 𝐾_d_^𝑡𝑑𝑃𝐶𝑃−𝑏^is the dissociation constant of the tdPCP-b protein produced in *E. coli*, which is not sialylated, 𝐾_d_^𝑠𝑖𝑎𝑙𝑜𝑝𝑟𝑜𝑡𝑒𝑖𝑛^ is the dissociation constant of a sialoprotein, 𝑙𝑛 is the natural logarithm, and ΔΔG is Gibbs free energy.

### Viral infection of Vero E6 Cells with SARS-CoV-2 and analysis

The experiments were carried out in a GLP-certified dedicated EU BSL3 (Bio-Safety Level 3) facility. Vero E6 cells(61) were seeded in a 96-wells plate (columns 1 to 11) in 100µl of M199 growth media (Gibco, cat. 41150087, supplemented with 5%v/v FBS) and allowed to adhere and grow overnight for 90% confluency (culture plate). The next day, a parallel plate was prepared (infection plate), holding both ACE2 granules and coronavirus virions (Delta or Omicron BA.1 variants). ACE2 and RNA highest tested concentration was concentrations were ∼0.2µM and ∼0.02M, respectively. All tested conditions were diluted 3-fold seven times in triplicates. The plate was prepared as follows: 52.5µl of ACE2 granules, or ACE2-only or RNA-only were added to row A of columns 1-3, 4-6, and 7-9, respectively. 52.5µl of infection media (M199, supplemented with 0.3% BSA) was added to columns 10-12, where untreated infected cells, untreated uninfected cells, and no cells controls would populate columns 10, 11, and 12, respectively. To rows B through H, 35µl of infection media was added. Using multichannel pipette, 17.5µl of the content of row B was transferred to row C, and the process repeated until row G. This comprised dilutions 0-7. 35µl of virus (∼2.5x10^5^ PFU/ml) was added to each well excepts uninfected columns (columns 11 and 12) which had received 35µl of infection media instead. Each well in the infection plate then held 70µl of infection media-virus-test article, and the infection plate was incubated for 1h at 37°C in a humidifier incubator. Following the incubation time, media was replaced with fresh media in the culture plate and the infection plate content was transferred to the culture plate. Infection was allowed to take place for 6h inside the incubator. Cell density was ∼2x10^4^ cells/well, with an MOI of ∼0.5.

Following the infection period, media was removed, and cells were washed once with PBS. Then 100µl of 4% formaldehyde in PBS was added to each well for fixation of cells. Plates were incubated at room temperature protected from light for 30min, and afterwards washed once with PBS. Cells were stored in 100µl fresh PBS at 4°C and stained within 2 days using AF488-labelled anti-spike antibody. Subsequently, plates were analyzed using CellInsight CX5 (Thermo Fisher).

### Influenza virus entry assay and analysis

Different sialogranules were tested on different days and results were normalized to the basal level of infected, untreated cells of the day of measurement (Figure 5C). 24h pre-infection, 2x10^4^ A549 cells(62) were seeded in two 96-wells plates in 100µl of DMEM growth media (Sigma, cat. D5796-500ML, supplemented with penicillin-streptomycin solution (IMBH, cat. L0022-100), FBS (IMBH, cat. S1400-500, and MEM NEAA solution (Sartorius, cat. 01-340-1B)) and allowed to adhere and grow overnight for 90% confluency. On the day of infection, two 96-well infection plates were prepared with influenza virions (A\H1N1\P.R.\8\34 strain) in the presence or absence of sialogranules or sialoprotein-only. The highest amount of sialoproteins was picomole, constituting a final concentration of 0.447mg/ml. All tested conditions were diluted 3-fold eight times. Sialogranules and sialoprotein-only were repeated in triplicates (LAMP1) or duplicates (GYPA, tdNXST1m, tdPCP), as were infected and untreated cells. The plates were prepared as follows: 52.5µl of sialogranules or sialoprotein-only were added row B. 35µl of infection media (growth media without FBS and with type IX-S trypsin (Sigma, cat. T0303-1G)) was added to subsequent rows (until row G) and rows B-D in another plate. Using a multichannel pipette, 17.5µl of the content of row B was transferred to row C, and the process repeated until row G. This comprised dilutions 0-5. 17.5µl was then transferred from row G to row B in the other plate and from there to rows C and then D. This comprised dilutions 6-8. 1µl of virus (1.8x10^5^ ∼PFU/ml) was added per 65µl of infection media and dispensed per well in columns 2-11 in both plates, bringing the final volume of infection media-virus-treatment (i.e. sialogranule or sialoprotein-only) to 100µl, and uninfected control received 100µl infection media without virus. Rows A and H and columns 1 and 12 in the infection plates were used as untreated, uninfected control. Columns 2 and 11 were used as untreated, infected control.

The infection step was carried out on two 96-well plates containing cells, as follows. Growth media was removed, and cells were washed twice with 50µl of PBSx1 to remove residual serum. Finally, the contents of the infection plates were transferred to the culture plates using a multi-channel pipettor, for an incubation period of 1h at 37 °C, 5% CO_2_. Following that, infection media was removed and replaced with fresh growth media, and the culture plates were returned to the same growing conditions. Infection was allowed to take place for 48h. Viral MOI was 0.01.

Following the period of infection, cells were detached and transferred to a U-bottom 96-well plate (Greiner, cat. 650101) in growth media. Cells were then centrifuged at 1500rpm, 4 °C for 5min. Media was removed without disturbing the cell pellet in the wells, and cells were resuspended and washed in 100µl FACS buffer (PBSx1, supplemented with 1%w/v BSA (Sigma, cat. A9418-50G)), then centrifuged again. The rinse step was repeated once more, before resuspending cells with 100µl CR6261 primary antibody solution (FACS buffer, supplemented with 1mg/µl of mouse-anti-HA antibody, freshly made). Plates were incubated for 1.5h, 4°C. At the end of the incubation period, cells were washed again before staining with 100µl secondary antibody solution (FACS buffer, supplemented with 1:200 AF488-labelled donkey-anti-mouse antibody (ENCO, cat. 715-545-151) and 1:1000 dilution of live/dead stain (Thermo Fisher, cat. 65-0865-14)). Plates were then incubated for 1h, 4°C protected from light. Finally, cells were washed twice more, and resuspended in 200µl of FACS buffer.

Finally, cells were taken for flowcytometry analysis (Fortessa-II). The flow cytometer was calibrated to compensate for the different labels in the system using compensation beads as follows: for FITC and live/stain dye, dedicated beads (Thermo Fisher, cat. 01-111142) were used as follows: 1µl of beads and 1µl of antibody with the same label were added to 1ml of FACS buffer. Beads were then centrifuged and 800µl were removed before resuspension. The same process was used for unlabeled beads. For mCherry compensation, we used 1 drop in 1ml of dedicated beads (Thermo Fisher, cat. A54743) in 1ml FACS buffer. Analysis of results was performed using custom MATLAB code. Matlab’s fit function was used to estimate the IC_50_ value of wells treated with the test articles (Supplementary Figure S10), using the following model (equation #5):

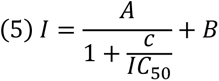

Where 𝐴 is the amplitude of response [0%,100%], 𝐵 is the intercept for the dependent variable 𝐼, 𝐼𝐶_50_ is the half-maximal inhibitory concentration, 𝑐 is the independent variable (test article concentration), and 𝐼 is the percent of infected cells in a sample. Test article concentration was used as mg/ml, and therefore the 𝐼𝐶_50_ units are the same. The fit guess was 𝐴=maximal mean percent of infected cells in the untreated uninfected samples; 𝐵=𝐴; 𝐼𝐶_50_=0.5.

### Numeric model for IFV virion flux measurement

We developed a simulation model to study a disrupted diffusion process of particles in a 3D bounded space with obstacles, incorporating particle attachment, hit counts, and particle generation mechanisms. The primary goal was to analyze the flux of particles hitting a cell layer, i.e., “the measurement wall”, arbitrarily designated as the x = 0 plane.

Model assumptions include (1) 3D bounded space with limits at [𝐿_𝑥_, 𝐿_𝑦_, 𝐿_𝑧_]; (2) particles undergo Brownian motion, characterized by a diffusion coefficient 𝐷; (3) virion flux is measured on the plane of 𝑥 = 0; (4) the space contains 𝐾 spherical obstacles which simulate a phase-separated condensate containing decoy receptors; and (5) upon hitting an obstacle, a virion becomes immobilized for a specified number of time steps 𝑡_𝑎𝑡𝑡𝑎𝑐ℎ_ before being released in a random direction.

The overall number of virion particles remained the same, and whenever a virion reached x = 0, a new virion was generated randomly in the 3D space. At the start of the simulation, 𝑁 particles and 𝐾 obstacles are initialized at random positions withing the bounded space. At each simulation step, particle positions are updated based on a diffusion process (equation #6):

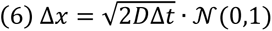

Where 𝒩(0,1) represents a number randomly generated from a standard normal distribution.

Particles are reflected back into the bounded space if they hit a boundary, as follows (equation #7):

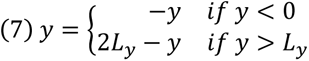

Similar reflection is applied for the 𝑧 dimension.

For the 𝑥 dimension the following applies (equation #8):

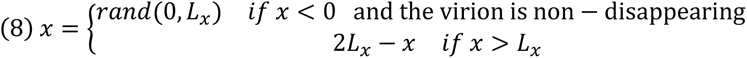

Where 𝑟𝑎𝑛𝑑(0, 𝐿_𝑥_) indicates a number randomly generated from a uniform distribution in the range [0, 𝐿_𝑥_].The number of particles hitting the measurement wall is recorded, and the flux rate is calculated at the end of the simulation as (equation #9), where *S* is the total number of simulation steps (10,000):

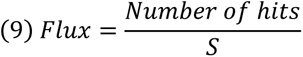

### Priming model

In order to model virus priming in a controlled cell culture setting, we have set up the following system of equations for *n* priming steps:

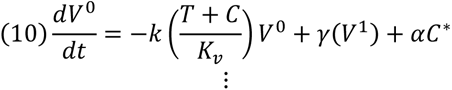

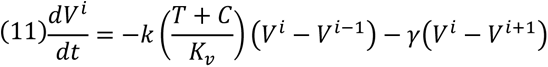

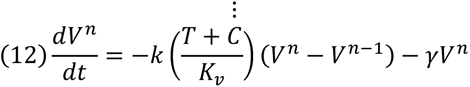

Where *V^0^*, *V^i^*,and *V^n^*correspond to the unprimed, i^th^-step, and n^th^-step primed virus concentration respectively. *T* and *C* correspond to the therapeutic and uninfected cell concentrations respectively. *C\**corresponds to infected cell concentration which creates new unprimed virus particles at a rate *α*. *K_v_* is the virus binding affinity to ACE2 (either as a therapeutic or on the cell), *k* is the rate at which the virus binds ACE2 leading to an increase in the “primed” state of the virion, and γ corresponds to a spontaneous reversion of the virion to a lower “primed” state. The priming equations are then complemented by the following equations:

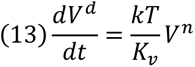

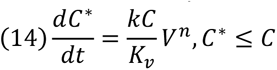

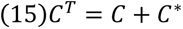

where *V^d^* corresponds to the concentration of deactivated virus particles as a result of the binding of a fully primed virion to a decoy therapeutic, the rate of creation of infected cells *C\** is provided by the probability for a fully primed virion (*V^n^*) to interact with an uninfected cell (*C*), and d *C^T^* is a constant corresponding to the total cell concentration in the simulation (1e6 in our settings). Simulation was implemented in Matlab 2021b (The MathWorks). See code availability statement for more detail

### Numeric model for SARS-CoV-2 virion flux measurement

We developed a simulation model to study a disrupted diffusion process of particles in a 3D bounded space with obstacles, incorporating particle attachment, hit counts, and particle generation mechanisms. The primary goal was to analyze the flux of particles hitting a cell layer, i.e., “the measurement wall”, arbitrarily designated as the x = 0 plane.

Model assumptions include (1) 3D bounded space with limits at [𝐿_𝑥_, 𝐿_𝑦_, 𝐿_𝑧_]; (2) particles undergo Brownian motion, characterized by a diffusion coefficient 𝐷; (3) virion flux is measured on the plane of 𝑥 = 0; (4) the space contains 𝐾 spherical obstacles which simulate a phase-separated condensate containing decoy receptors; and (5) upon hitting an obstacle, a virion becomes immobilized for a specified number of time steps 𝑡_𝑎𝑡𝑡𝑎𝑐ℎ_ before being released in a random direction.

Particles were assigned a hit count 𝐻 (representing priming) – when a particle hits an obstacle or the measurement wall, its hit count increases by 1. If a particle with 𝐻 − 1 hits reaches 𝐻 hits upon hitting an obstacle, it disappears. If a particle with 𝐻 − 1 hits reaches 𝐻 upon hitting the measurement wall, it disappears and 𝑃 new particles are generated at random positions withing the bounded space (simulating successful infection).

At the start of the simulation, 𝑁 particles and 𝐾 obstacles are initialized at random positions withing the bounded space. At each simulation step, particle positions are updated based on a diffusion process (equation #6):

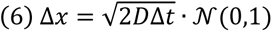

Where 𝒩(0,1) represents a number randomly generated from a standard normal distribution.

Particles are reflected back into the bounded space if they hit a boundary, as follows (equation #7):

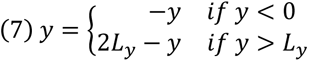

Similar reflection is applied for the 𝑧 dimension.

For the 𝑥 dimension the following applies (equation #8):

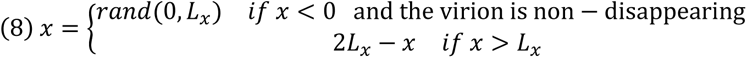

Where 𝑟𝑎𝑛𝑑(0, 𝐿_𝑥_) indicates a number randomly generated from a uniform distribution in the range [0, 𝐿_𝑥_]. Particles hitting the measurement wall (*x* = 0) have their hit counts increased by 1. If a particle’s hit count reaches 𝐻 at the measurement wall, it disappears and 𝑃 new particles are generated in the space. For particles hitting obstacles, their hit counts are increased by 1. If a particle’s hit count reaches 𝐻 at an obstacle it disappears. Otherwise, it attaches to the obstacle for 𝑡_𝑎𝑡𝑡𝑎𝑐ℎ_ time steps. After this period, it is released in a random direction.

The number of particles hitting the measurement wall is recorded, and the flux rate is calculated at the end of the simulation as (equation #9), where *S* is the total number of simulation steps (10,000):

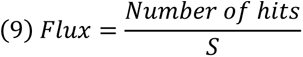

### Statistics and reproducibility

The statistical analysis was calculated using a custom Matlab code. For in vitro image analysis experiment, the data for each protein was obtained from several hundred granules. For distribution comparison, a standard two-sided student t-test was used.

All infection experiments were carried out in biological triplicates. ACE2 and LAMP1 were performed in triplicates, while GYPA and tdNXST1 were performed in duplicates.

No statistical method was used to predetermine sample size. No data were excluded from the analyses. The experiments were not randomized. The investigators were not blinded to allocation during experiments and outcome assessment.

## Data availability statement

The data supporting the findings of this study is publicly available at Zenodo under accession code https://zenodo.org/doi/10.5281/zenodo.12789664.

## Code availability statement

The code, trained models, and processed datasets are publicly available at https://zenodo.org/doi/10.5281/zenodo.12789704 and at 10.6084/m9.figshare.26830774.

## Author Contributions

**OW** designed and cloned the constructs for 𝞓ACE2, ACE2NB, RBD-sfGFP-His, LAMP1, GYPA, Fetuin, the NXST variants, as well as designed and carried out the experiments and analysis for most of the data. **NG** and RA conceived the priming model. **NG** coded the priming model and the virus infection simulations and inferred the protein structures using AlphaFold. **OW** and **SG** designed the construct for wild-type ACE2. **SG** designed and cloned tdPCPm, cloned NXST1b, and guided the microscopy experiments and image analysis. **RA** supervised the study. **OW** and **RA** wrote the manuscript.

## Acknowledgements

We thank the Life Sciences and Engineering (LS&E) Infrastructure Center at the Technion (Dr. Nitsan Dahan and Dr. Yael Lupu-Haber from the microscopy core facility) for their excellent technical assistance with confocal microscopy imaging. We thank Dr. Yotam Bar-on and Dr. Orly Kladnitsky from the Faculty of Medicine at the Technion for the IFV strain and the materials required for its viral entry assay. We would like to express our gratitude to Amir Grau from the Cytometry Center at the Rappaport Faculty of Medicine Biomedical Core Facility (BCF) for assisting us with the flow cytometry technology. We would like to thank Benevira, Inc. CEO Patricia Kitchen for funding the Omicron and Delta entry assay and for Viroclinics Inc. for executing it in GLP conditions. This work was greatly supported by the European Union’s Horizon 2020 Research and Innovation Programme under grant agreements no. 851615 and no. 851065.

## Competing Interests

The authors declare the following competing financial interest(s): OW, NG, SG, and RA are inventors on US Provisional Patent Application No. 63/187969 concerning some of the described technologies. RA is an inventor on US Patent Application 2021/0095296 A1.

**Supplementary Movie S1 – tdPCPb is not affected by the addition of SNA.** Blue – slncRNA. Red – tdPCPb.

**Supplementary Movie S2 – tdPCPm transitions from granular shapes into a liquid-like dense cloud of SNA and sialoproteins, with slncRNA at the periphery.** Blue – slncRNA. Green – SNA. Red – tdPCPm.

**Supplementary Movie S3 – 3D rendering of a LAMP1 granule.** slncRNA (blue) is clustered at the periphery of an SNA (green)-LAMP1 (red) overlap.

---

**Supplementary Figure S1.**
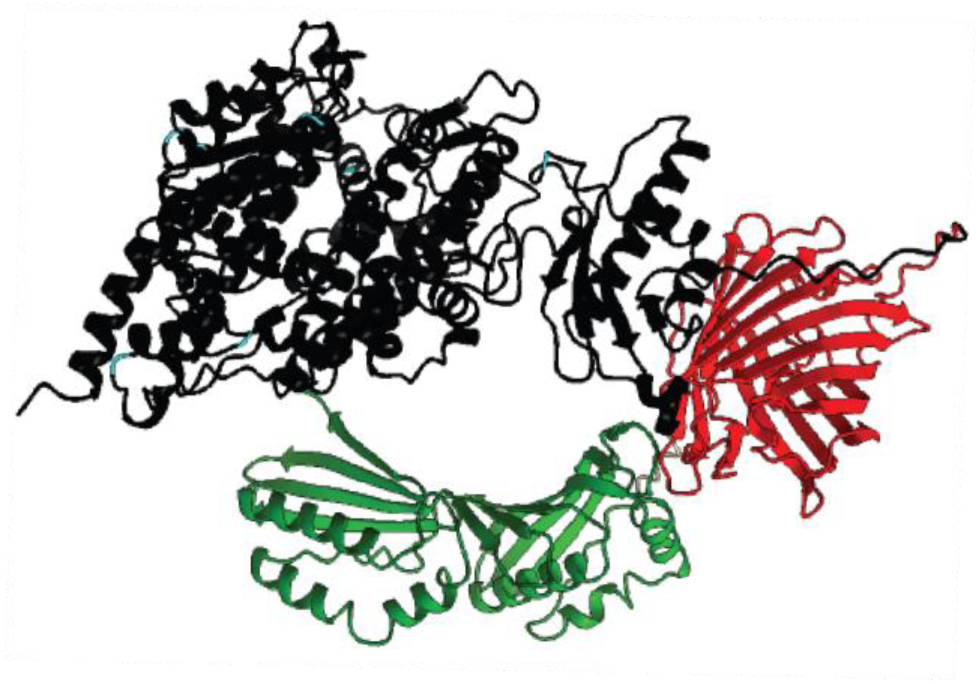
AlphaFold prediction of ACE2-mCherry-tdPCP structure. Black – ACE2. Red – mCherry. Green – tdPCP. Light blue markings – asparagine residues (part of the N-X-S/T motif).

**Supplementary Figure S2.**
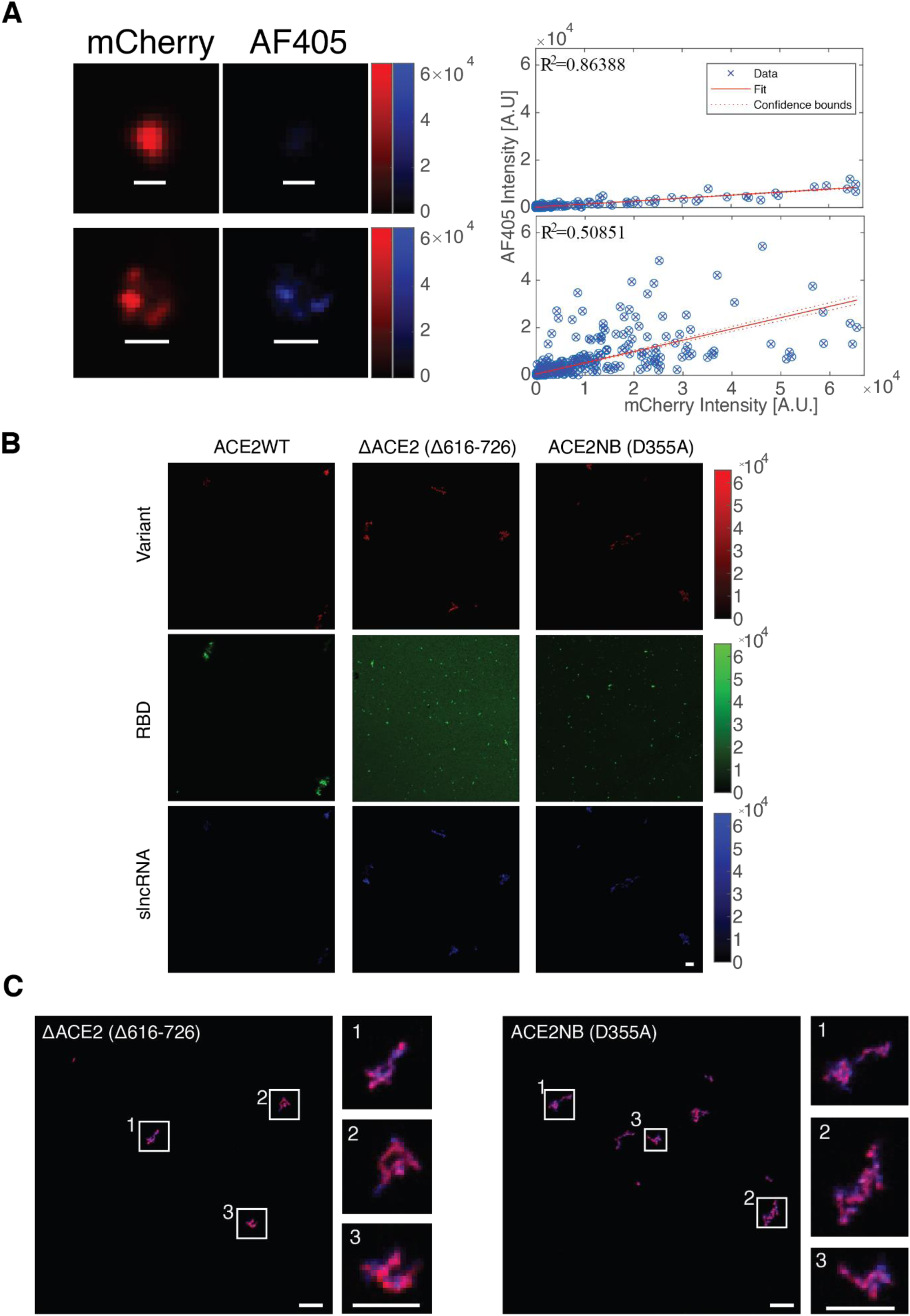
ACE2 (wild-type and mutants) interactions with slncRNA and RBD. (**A)** Representative correlations between mCherry and AF405 signal intensities for the two events in Figure 1B. **(B)** Individual channels of the ACE2 granules and RBD shown in Figure 1D, 1H (top left) and 1I (top left). **(C)** Granules formed by the ACE2 mutants.

**Supplementary Figure S3.**
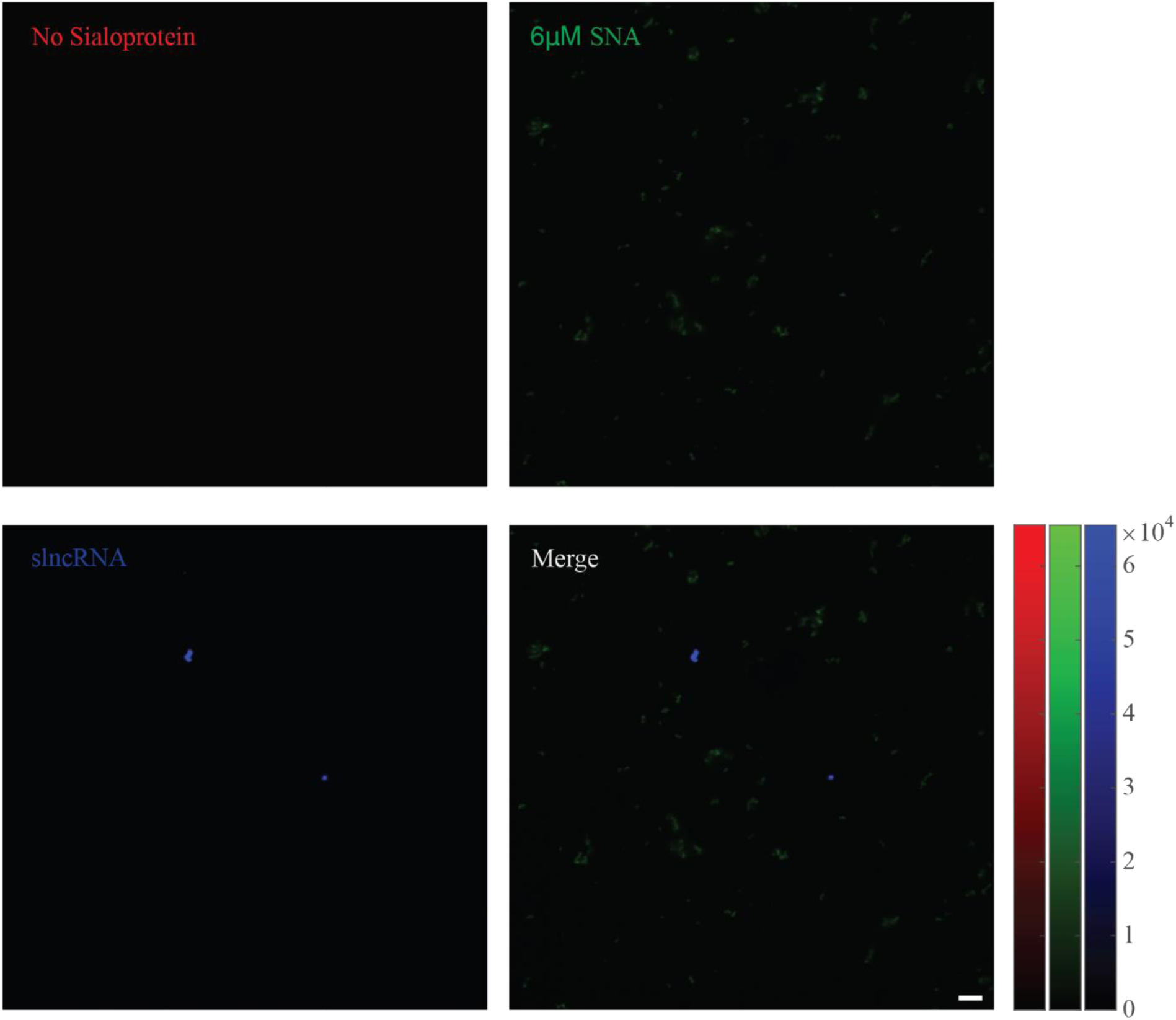
slncRNA and SNA do not interact without a sialoprotein. (Top left) mCherry channel (sialoprotein). (Top right) FITC channel (SNA). (Bottom left) AF405 channel (slncRNA). (Bottom right) Merged channels. Scalebar: 5µm.

**Supplementary Figure S4.**
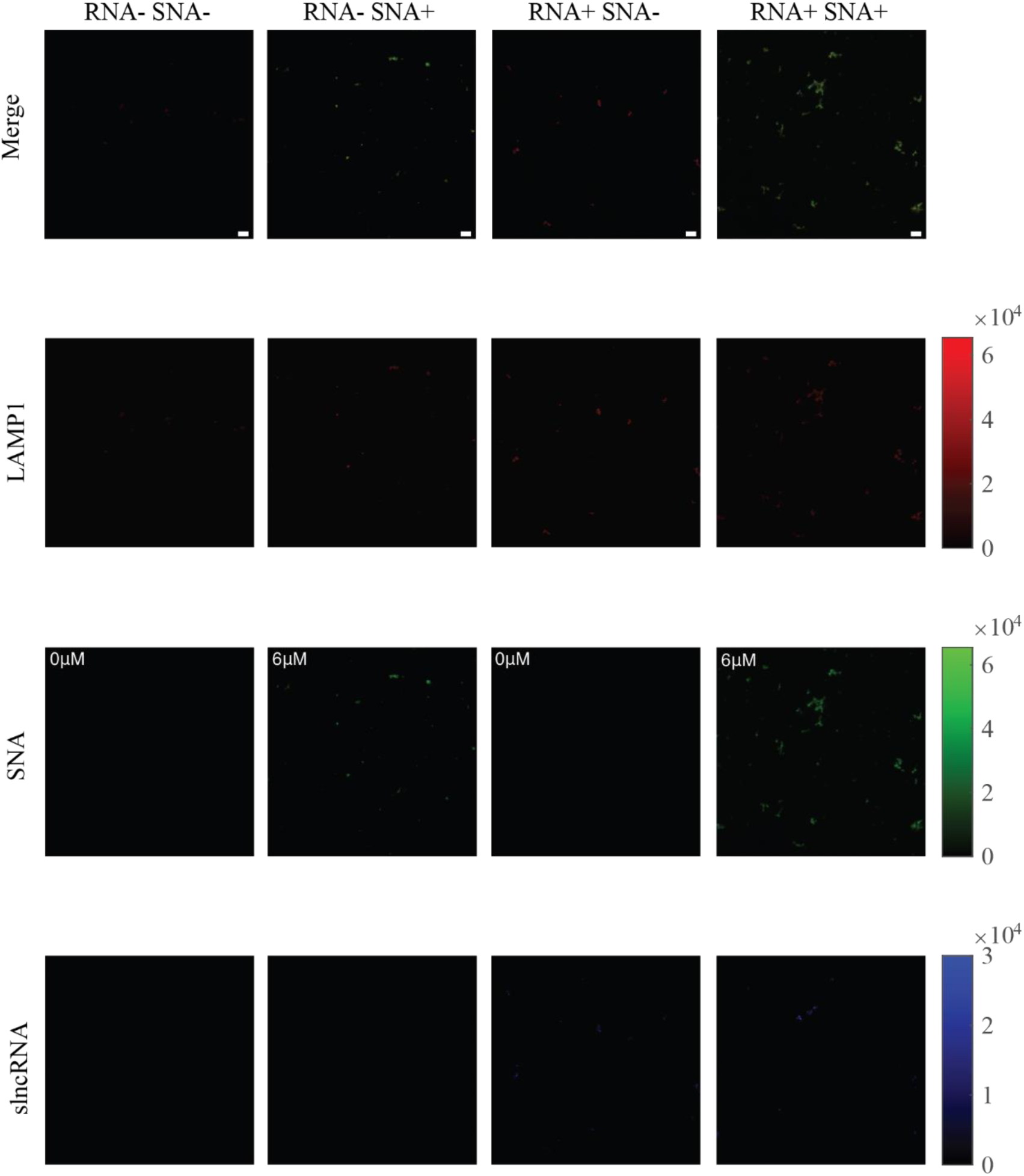
slncRNA presence enhances SNA-sialoprotein interactions. slncRNA presence results in larger structures that form once SNA is added. Additionally, the mCherry signal is strengthened by the presence of the slncRNA, as previously reported(11). Scalebar: 5µm.

**Supplementary Figure S5.**
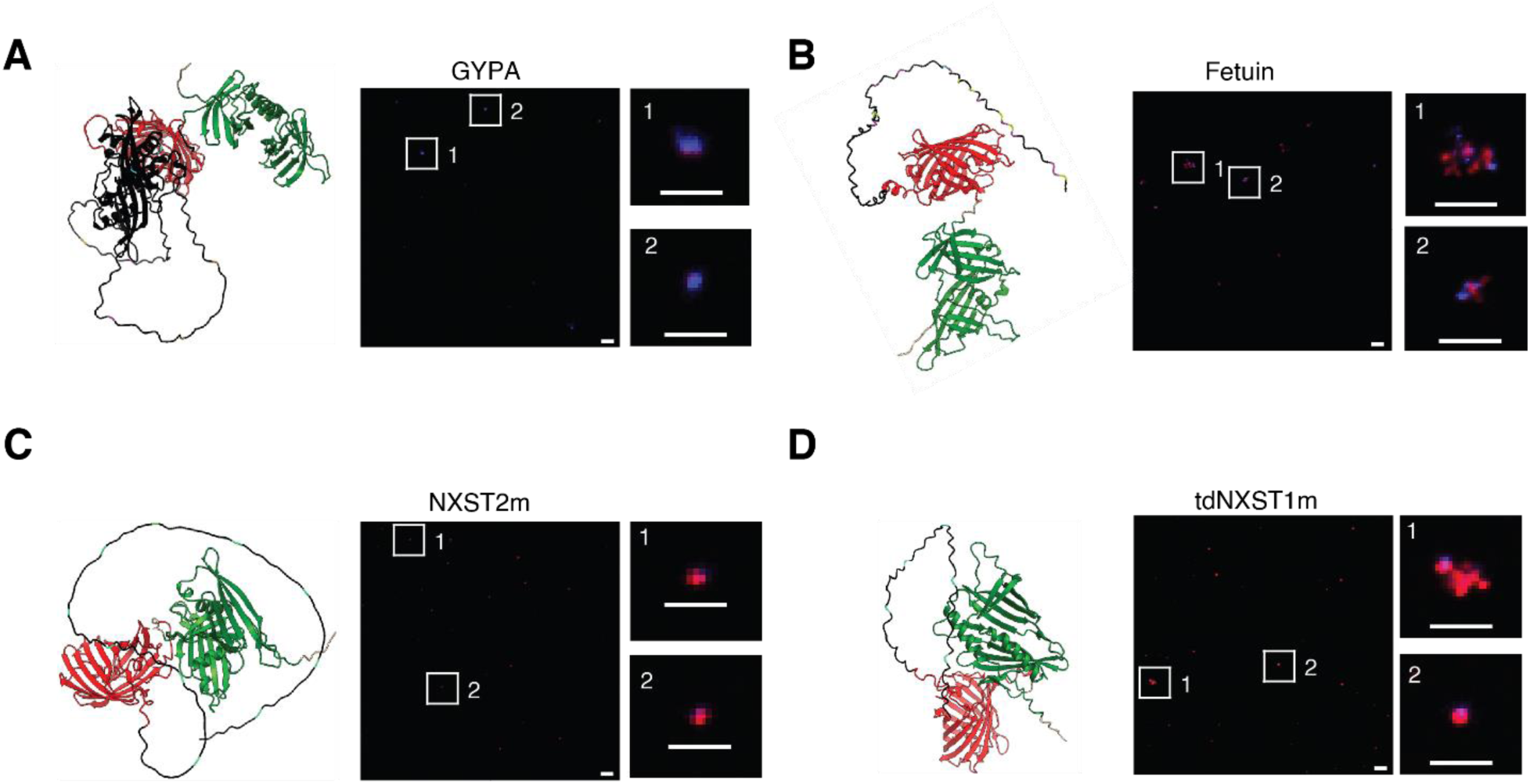
Sialoprotein candidates form granules in the presence of slncRNA. **(A)** GYPA. **(B)** Fetuin. **(C)** NXST2m. **(D)** tdNXST1m. Black, red, and green are sialoprotein, mCherry, and tdPCP, respectively. Putative sialylated sites are marked in light blue (asparagine residues, part of the N-X-S/T motif), yellow (serine), or magenta (threonine). Scalebar: 5µm (large FoV) and 2µm (events enlargement).

**Supplementary Figure S6.**
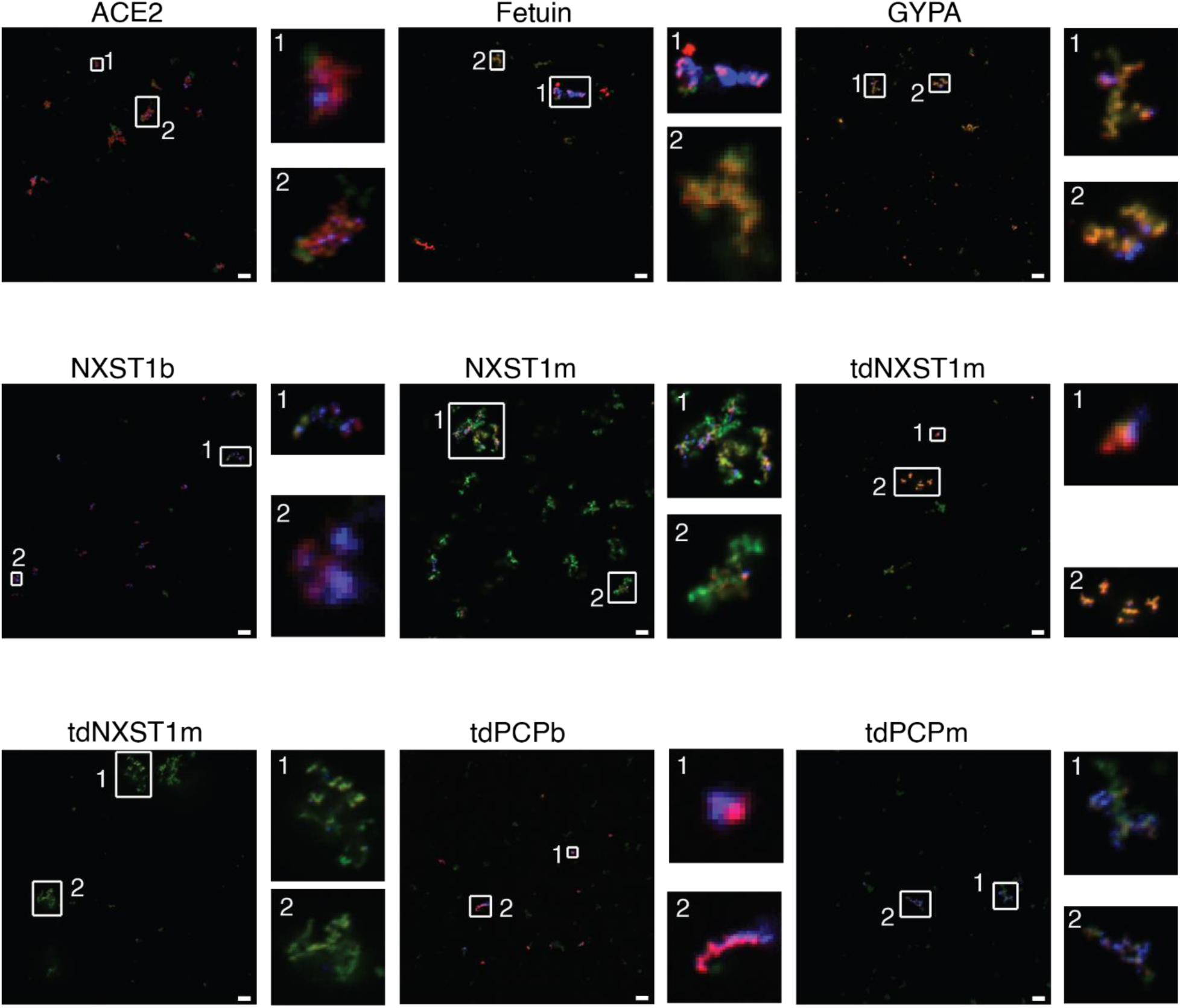
Addition of SNA results in the formation of biocondensates in nearly all sialoprotein candidates. Scalebar: 5µm.

**Supplementary Figure S7.**
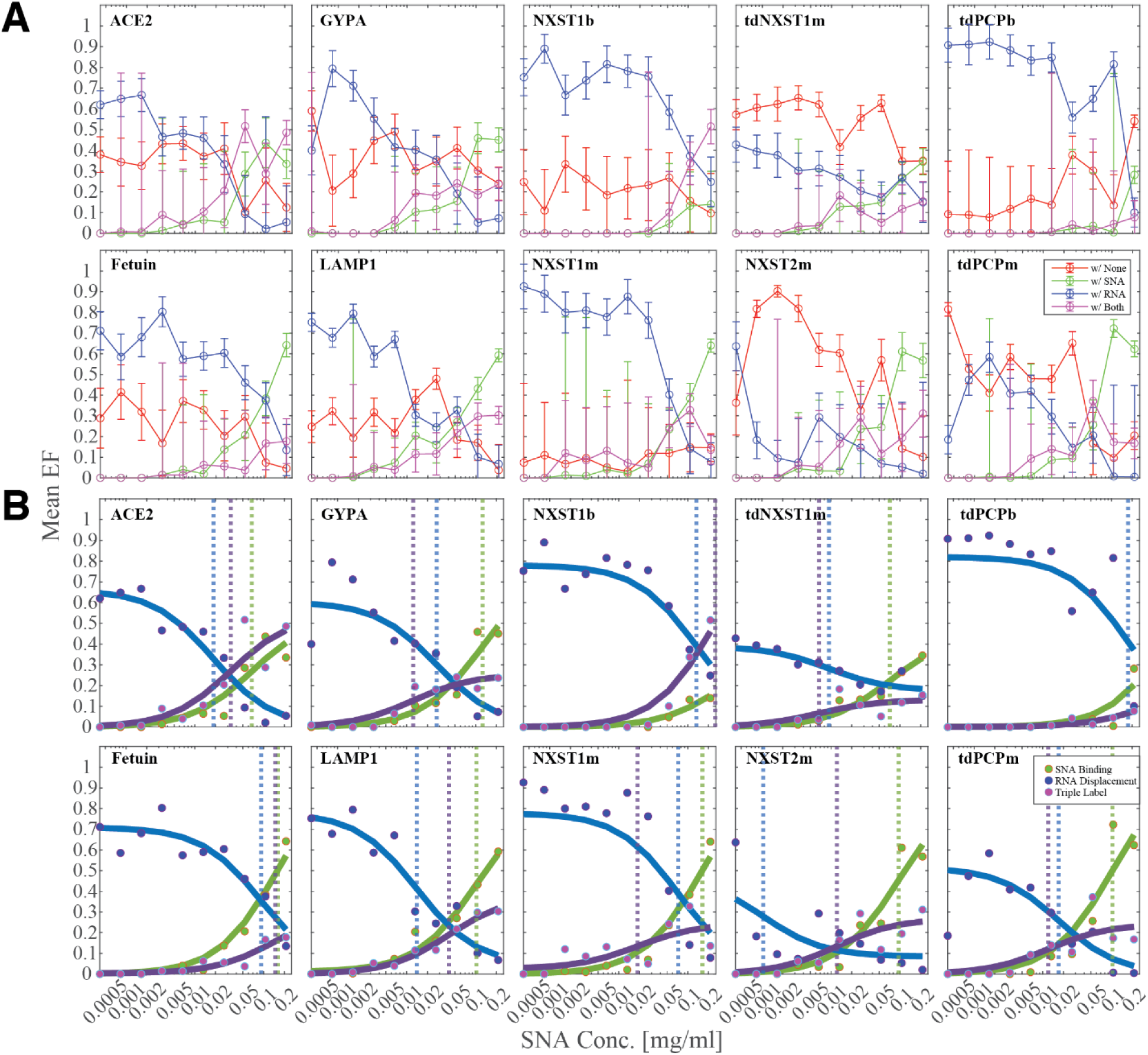
Mean event frequency (EF) across all sialoprotein candidates. **(A)** mCherry-positive events are classified as either colocalized with AF405 (RNA, blue), FITC (SNA, green), both (triple label, magenta), or neither (red) (see Methods and Figure 4c). As SNA concentration increased, colocalization with SNA and triple labelling increased while colocalization with RNA decreased. **(B)** Fitted curves of the data in Supplementary Figure7A. Dashed vertical lines mark the *K_d_* value for each fit line.

**Supplementary Figure S8.**
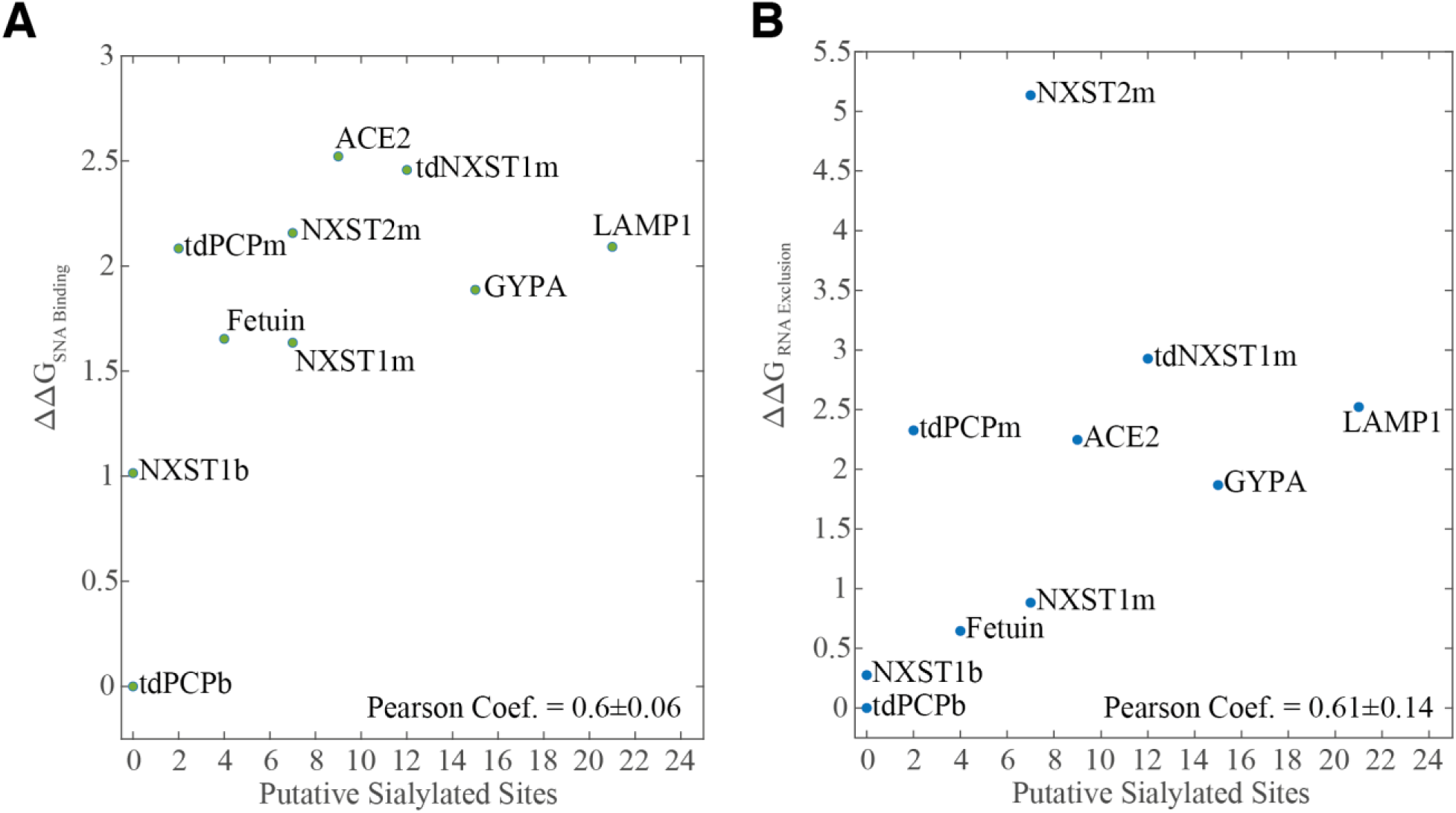
Gibbs’ free energy (𝞓𝞓G) for SNA binding and RNA displacement is dependent on the number of putative sialylated sites. A strong pearson correlation is demonstrated between the number of putative sialylated sites and either the accumulation of SNA signal **(A)** or reduction in RNA signal **(B)**. The higher the number of putative sialylated sites, the more the two processes occur.

**Supplementary Figure S9.**
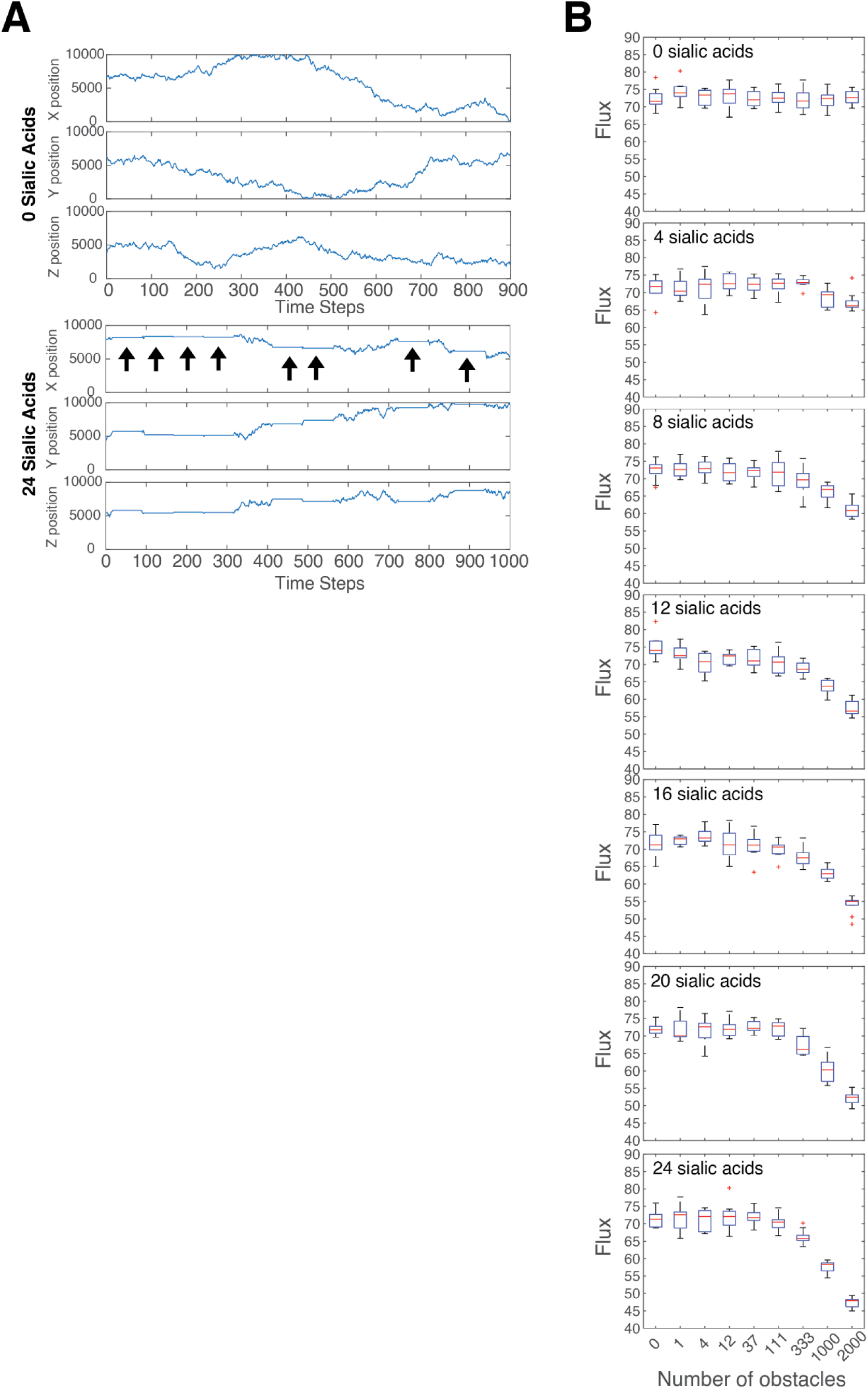
Obstacles that interact with IFV prolong viral diffusion time and reduce the viral infection flux. **(A)** Representative trajectories in 3D space of a viral particle in the absence (top) or presence (bottom) of obstacles that it can interact with (i.e. sialogranules). Interactions with an obstacle (marked by arrows) result in a delay in progression towards the cell layer and increases the time it takes to infect a cell. **(B)** The viral infection flux decreases with increasing number of obstacles (x-axis) or stronger the viral-obstacle interactions (different boxplots clusters).

**Supplementary Figure S10.**
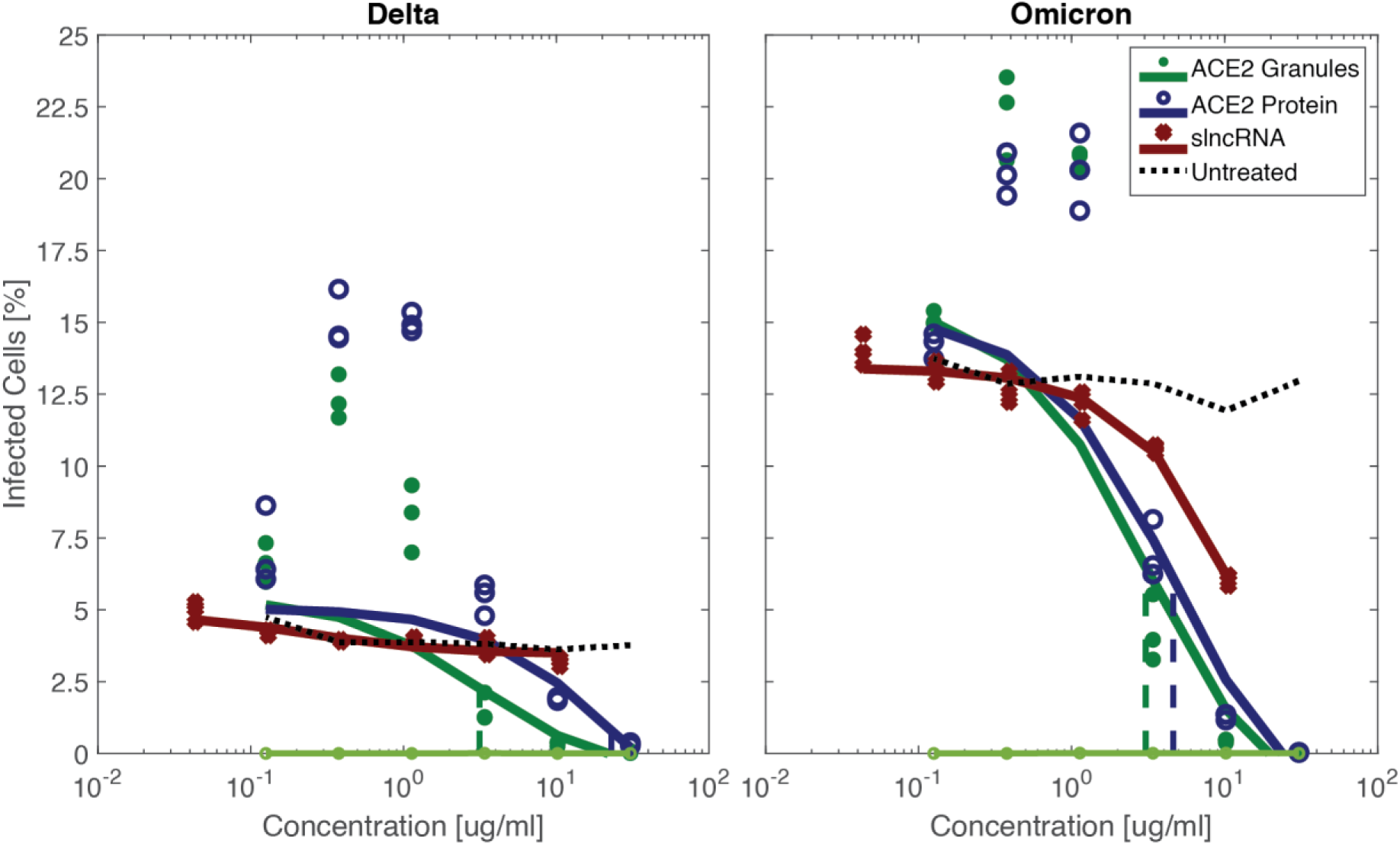
ACE2 decoy particles inhibit SARS2 infection at a lower IC_50_ value than an ACE2-only solution. Infection by either the Delta (left) or Omicron (right) variants was inhibited in Vero E6 cells using ACE2 decoy particles. This was achieved at a lower IC_50_ value for both variants.

**Table 1.**
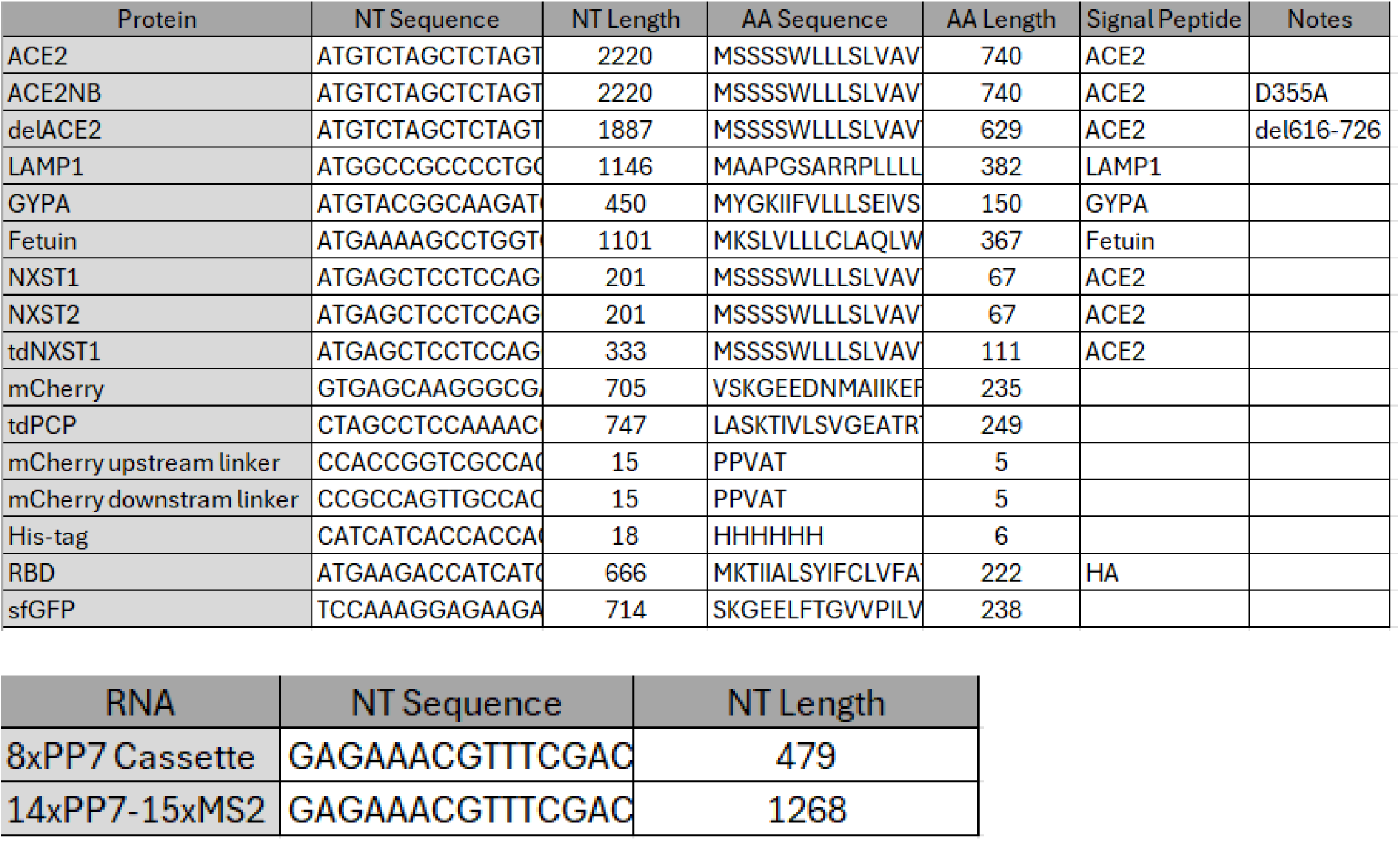
proteins and RNA sequences.

