## Supplementary Information for "Phase Separation-based Antiviral Decoy Particles as Basis for Programmable Broad-spectrum Therapeutics"

Supplementary Data

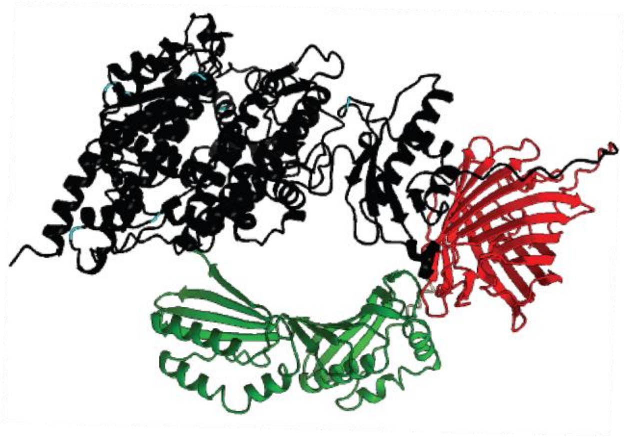

**Supplementary Figure S1 – AlphaFold prediction of ACE2-mCherry-tdPCP structure.** Black – ACE2. Red – mCherry. Green – tdPCP. Light blue markings – asparagine residues (part of the N-X-S/T motif).

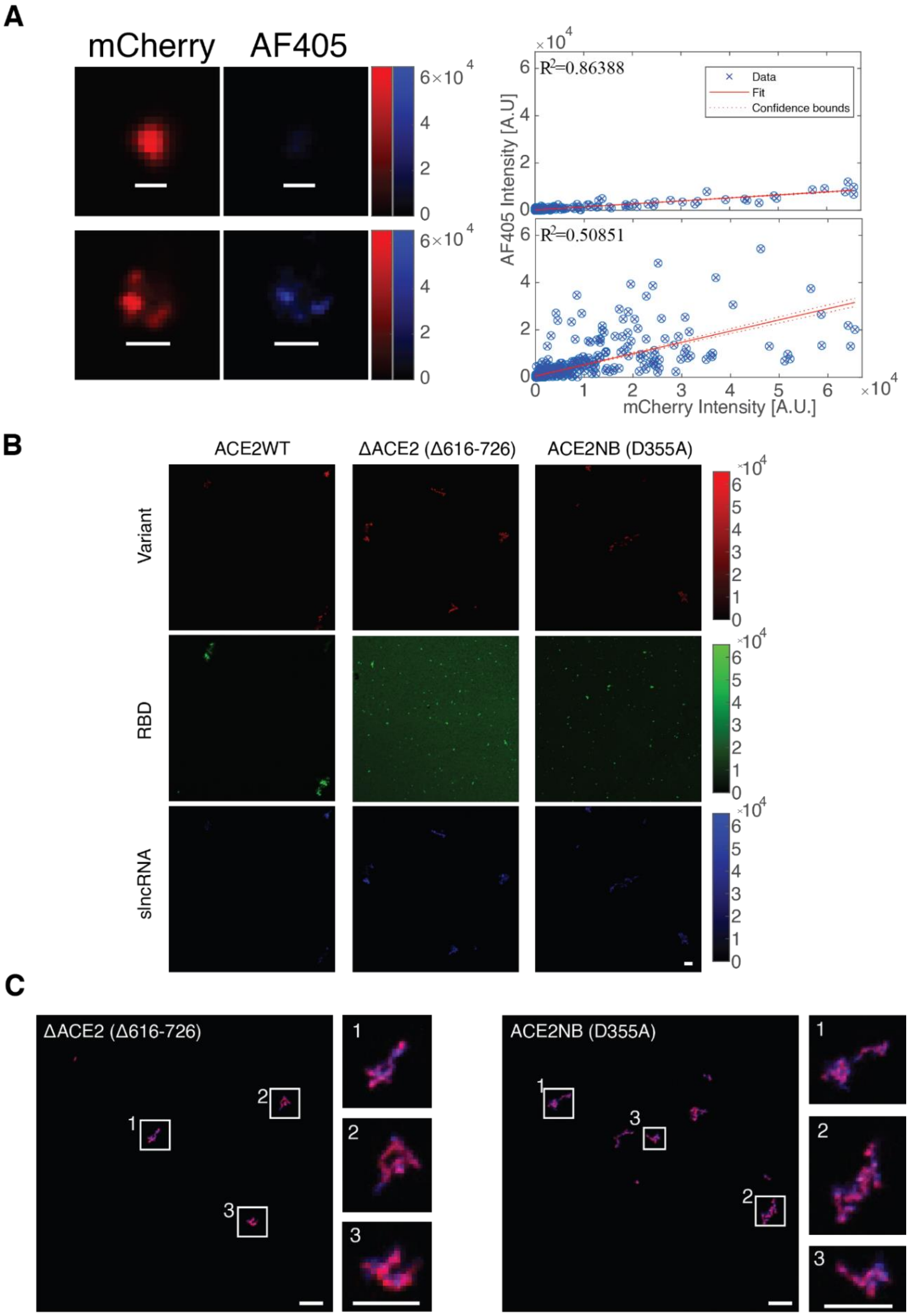


**Supplementary Figure S2 – ACE2 (wild-type and mutants) interactions with slncRNA and RBD.** (**A)** Representative correlations between mCherry and AF405 signal intensities for the two events in Figure 1B. **(B)** Individual channels of the ACE2 granules and RBD shown in Figure 1D, 1H (top left) and 1I (top left). **(C)** Granules formed by the ACE2 mutants.

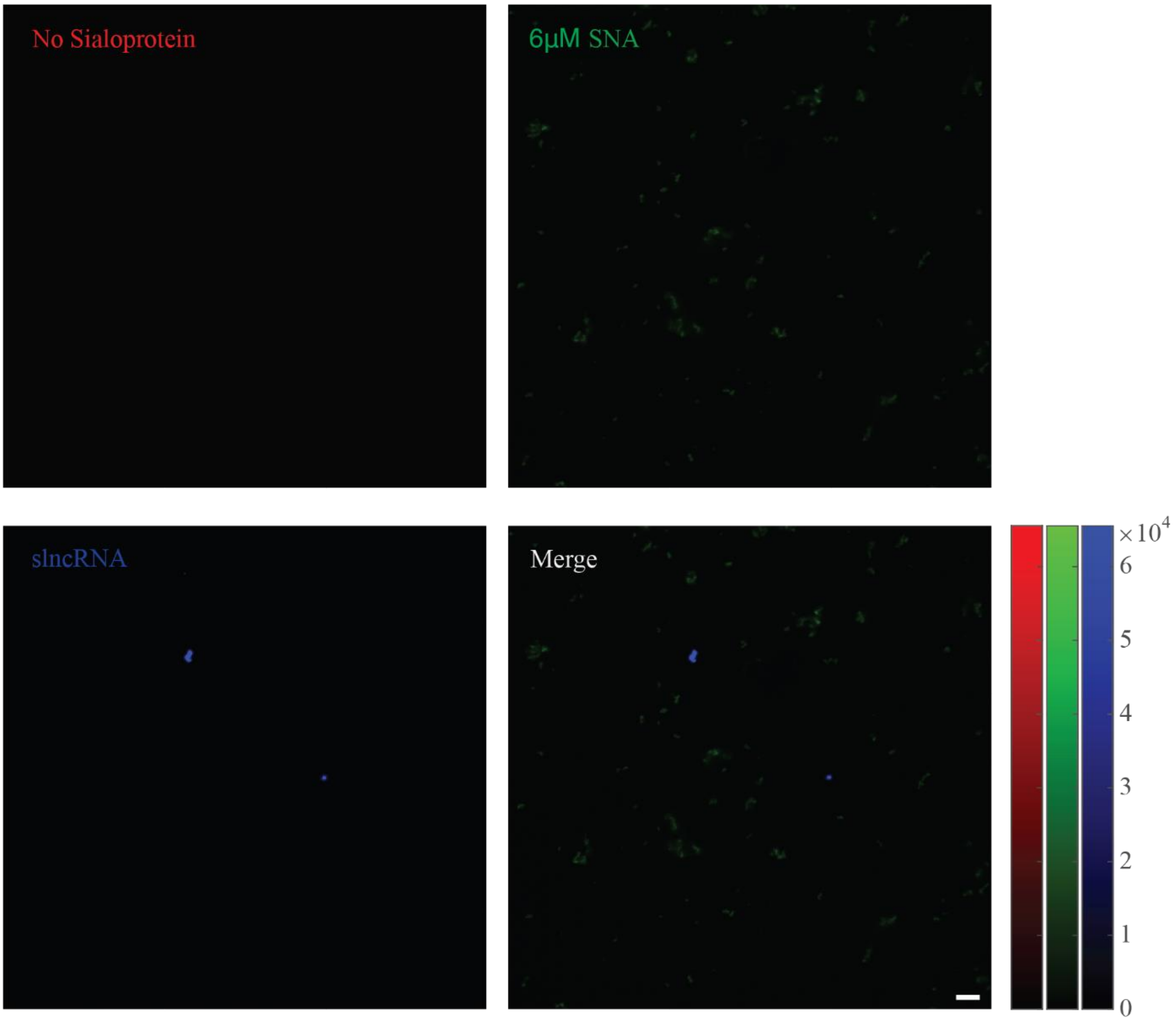


**Supplementary Figure S3 – slncRNA and SNA do not interact without a sialoprotein.** (Top left) mCherry channel (sialoprotein). (Top right) FITC channel (SNA). (Bottom left) AF405 channel (slncRNA). (Bottom right) Merged channels. Scalebar: 5μm.

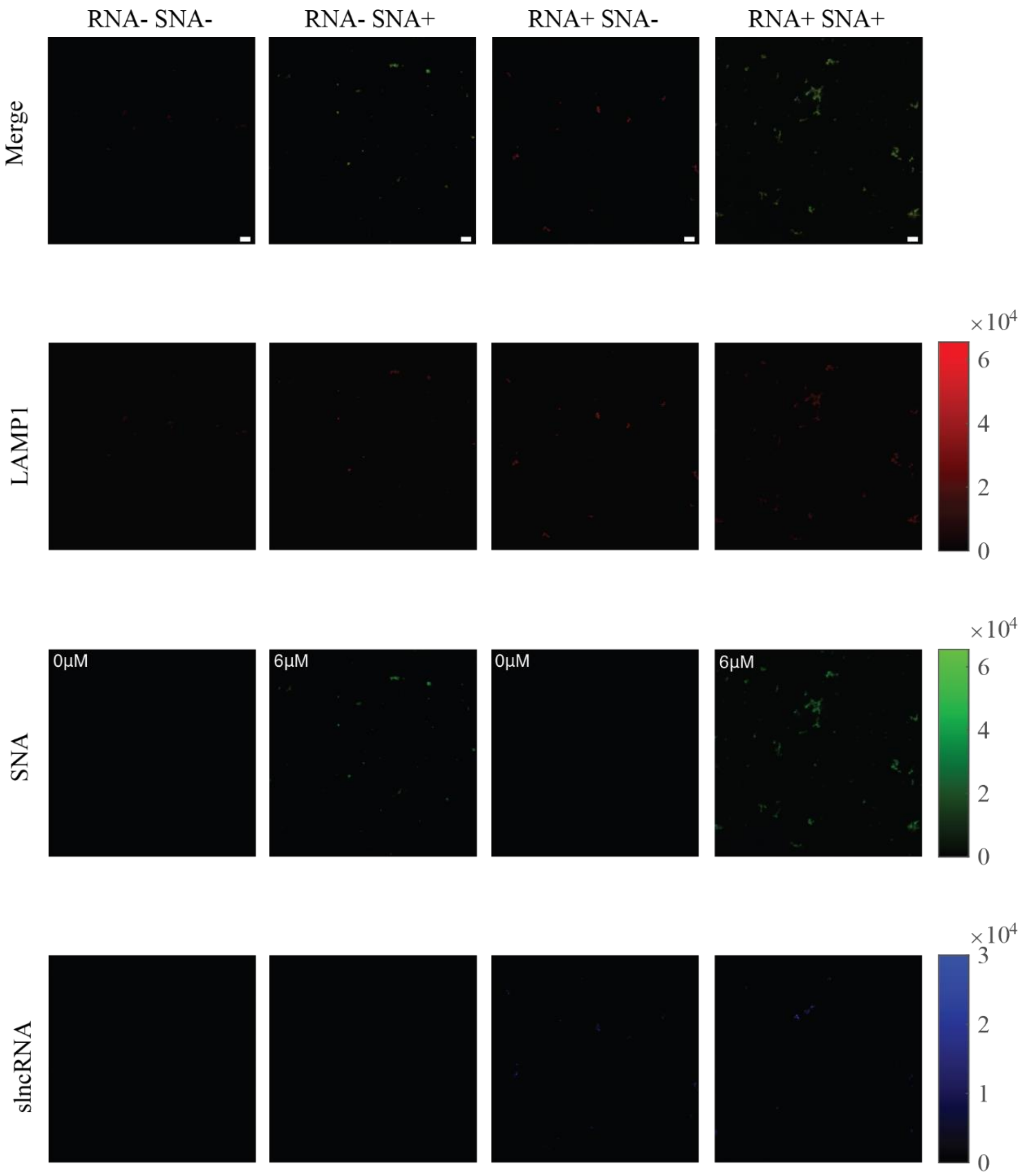


**Supplementary Figure S4 – slncRNA presence enhances SNA-sialoprotein interactions.** slncRNA presence results in larger structures that form once SNA is added. Additionally, the mCherry signal is strengthened by the presence of the slncRNA, as previously reported(11). Scalebar: 5µm.

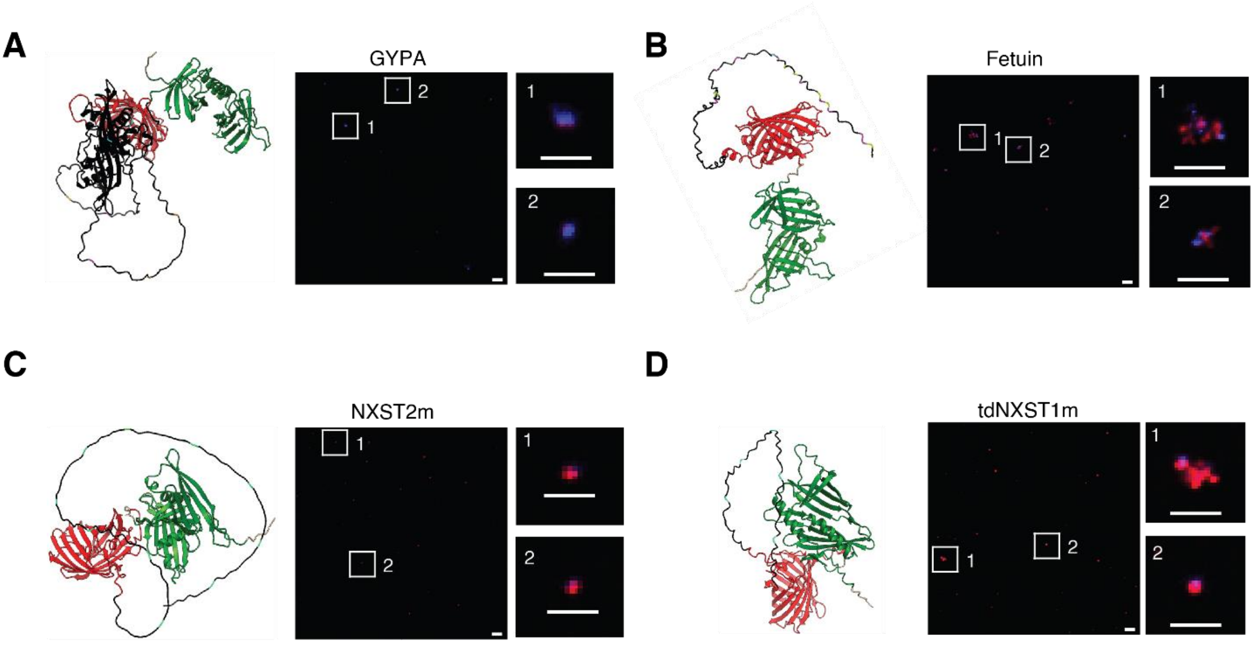


**Supplementary Figure S5 – Sialoprotein candidates form granules in the presence of slncRNA. (A)** GYPA. **(B)** Fetuin. **(C)** NXST2m. **(D)** tdNXST1m. Black, red, and green are sialoprotein, mCherry, and tdPCP, respectively. Putative sialylated sites are marked in light blue (asparagine residues, part of the N-X-S/T motif), yellow (serine), or magenta (threonine). Scalebar: 5µm (large FoV) and 2µm (events enlargement).

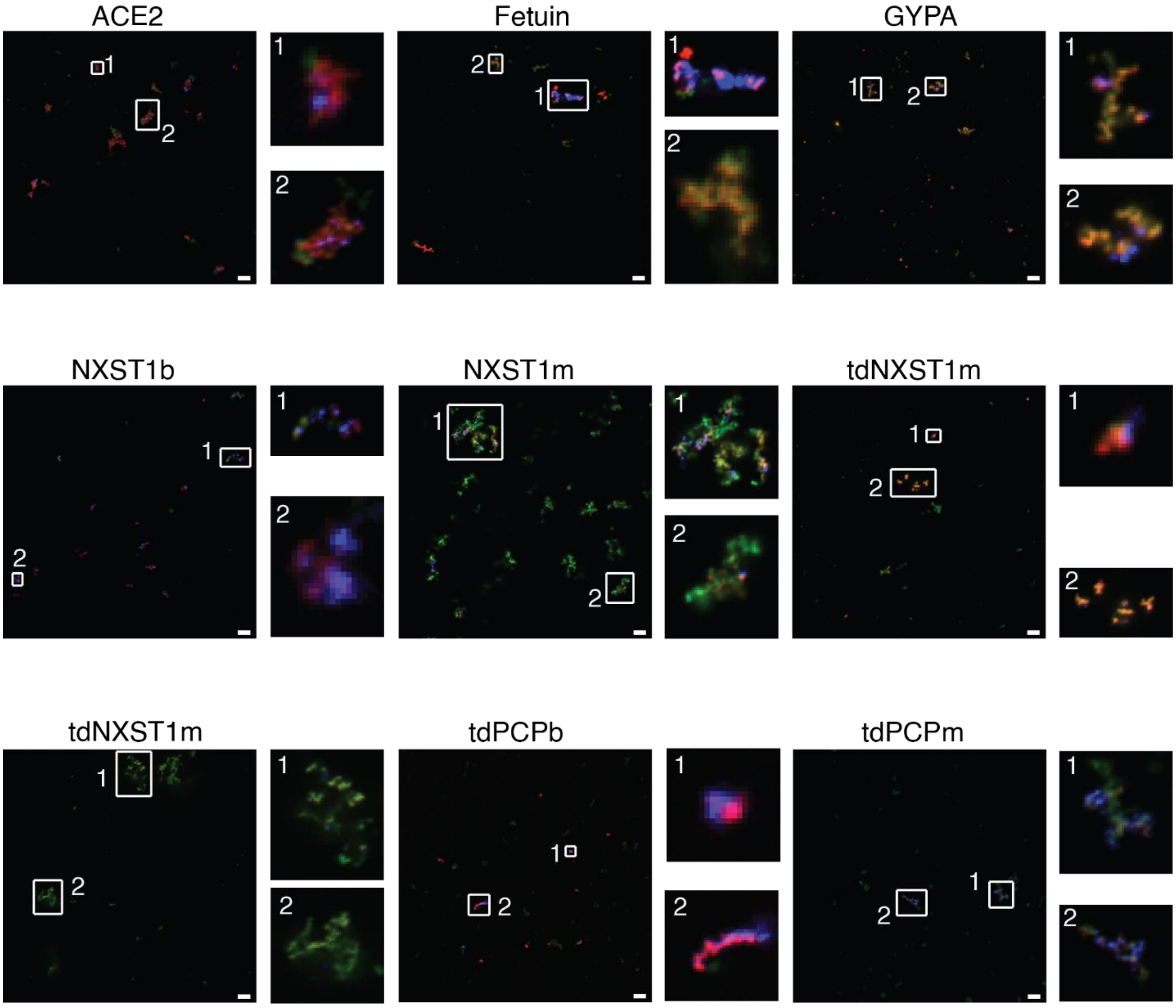


**Supplementary Figure S6 – Addition of SNA results in the formation of biocondensates in nearly all sialoprotein candidates.** Scalebar: 5μm.

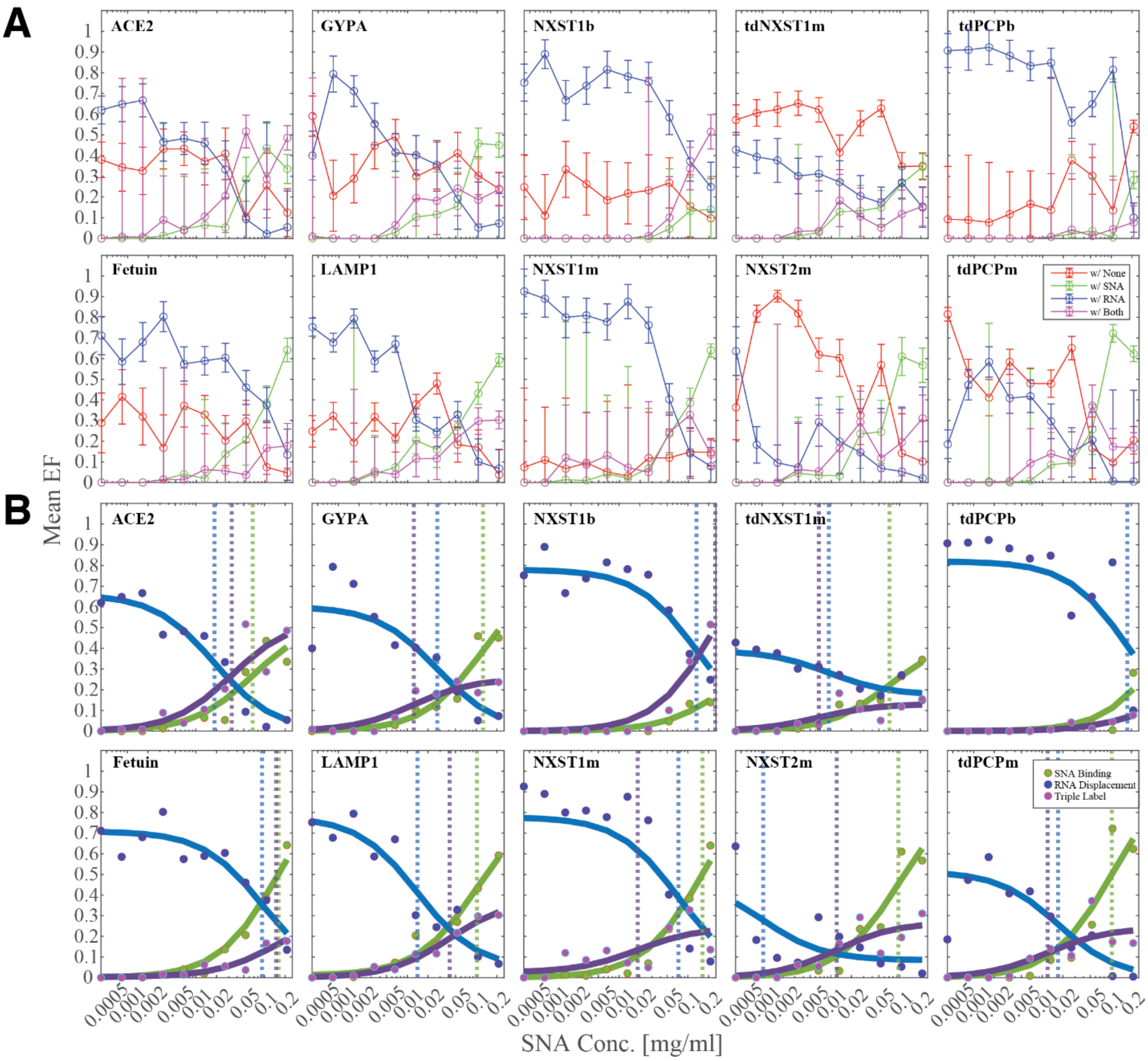


**Supplementary Figure S7 – Mean event frequency (EF) across all sialoprotein candidates. (A)** mCherry-positive events are classified as either colocalized with AF405 (RNA, blue), FITC (SNA, green), both (triple label, magenta), or neither (red) (see Methods and Figure 4c). As SNA concentration increased, colocalization with SNA and triple labelling increased while colocalization with RNA decreased. **(B)** Fitted curves of the data in Supplementary Figure7A. Dashed vertical lines mark the $K_d$ value for each fit line.

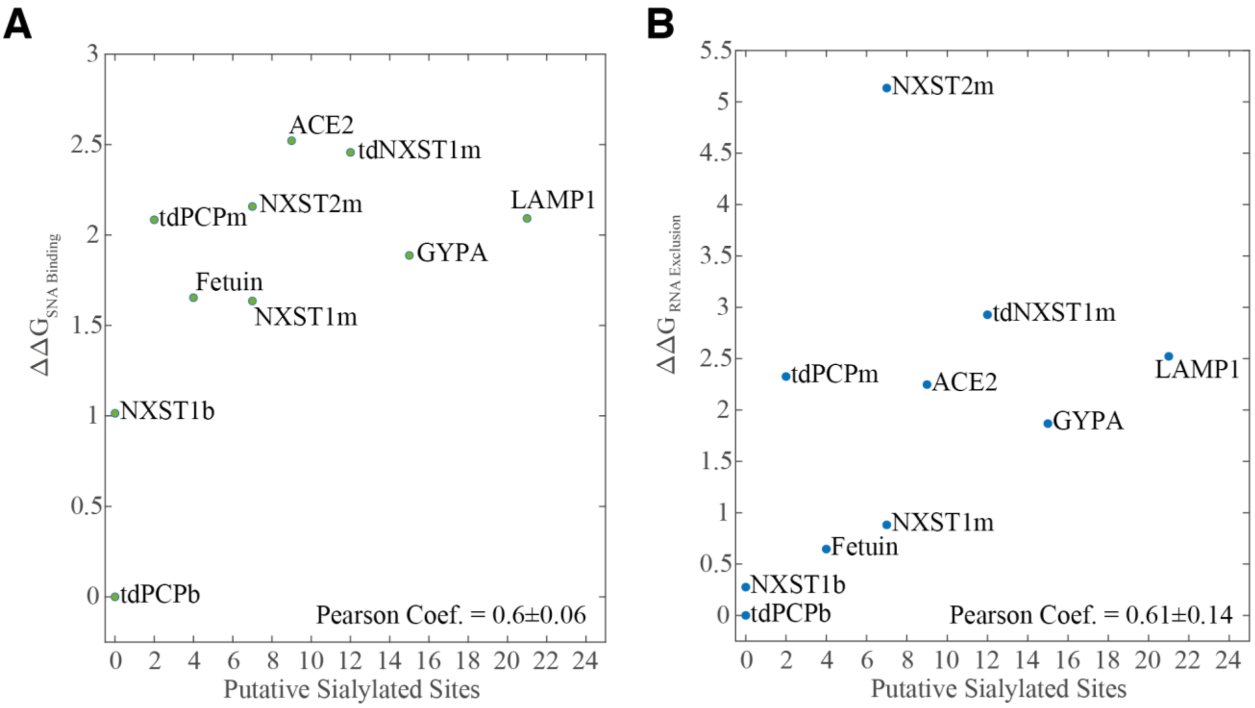


**Supplementary Figure S8 – Gibbs' free energy (*ΔΔ*G) for SNA binding and RNA displacement is dependent on the number of putative sialylated sites.** A strong pearson correlation is demonstrated between the number of putative sialylated sites and either the accumulation of SNA signal **(A)** or reduction in RNA signal **(B)**. The higher the number of putative sialylated sites, the more the two processes occur.

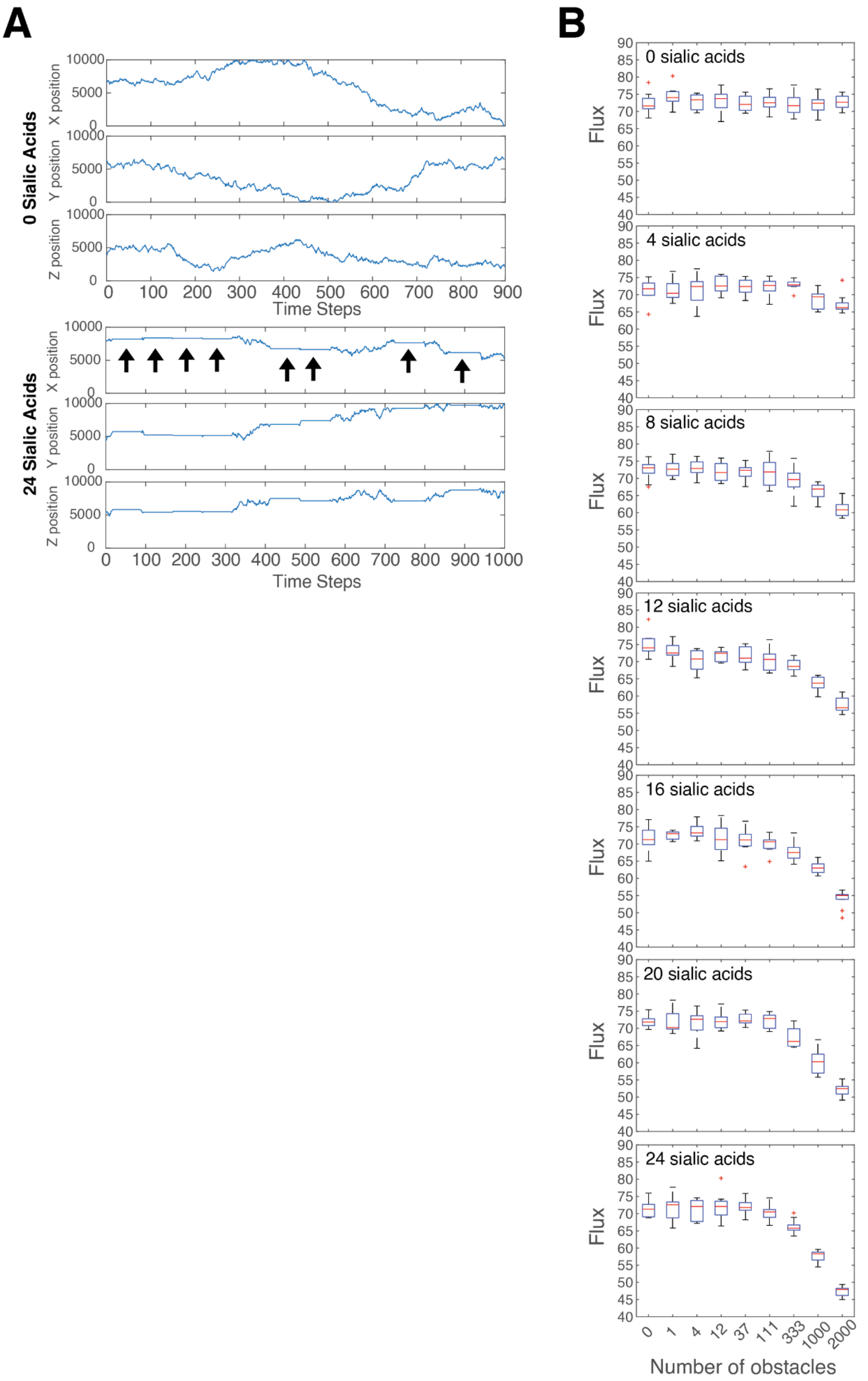


**Supplementary Figure S9 – Obstacles that interact with IFV prolong viral diffusion time and reduce the viral infection flux. (A)** Representative trajectories in 3D space of a viral particle in the absence (top) or presence (bottom) of obstacles that it can interact with (i.e. sialogranules). Interactions with an obstacle (marked by arrows) result in a delay in progression towards the cell layer and increases the time it takes to infect a cell. **(B)** The viral infection flux decreases with increasing number of obstacles (x-axis) or stronger the viral-obstacle interactions (different boxplots clusters).

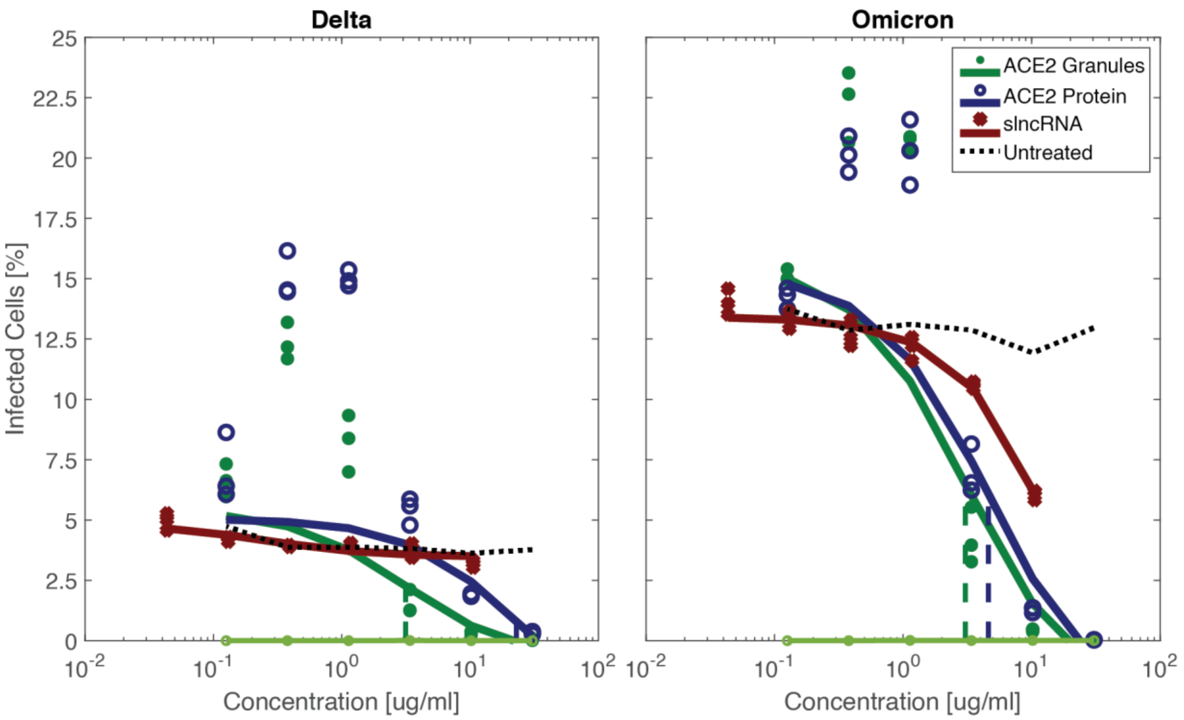


**Supplementary Figure S10 – ACE2 decoy particles inhibit SARS2 infection at a lower $IC_{50}$ value than an ACE2-only solution.** Infection by either the Delta (left) or Omicron (right) variants was inhibited in Vero E6 cells using ACE2 decoy particles. This was achieved at a lower $IC_{50}$ value for both variants.

**Supplementary Movie S1 – tdPCPb is not affected by the addition of SNA.** Blue – slncRNA. Red – tdPCPb.

**Supplementary Movie S2 – tdPCPm transitions from granular shapes into a liquid-like dense cloud of SNA and sialoproteins, with slncRNA at the periphery.** Blue – slncRNA. Green – SNA. Red – tdPCPm.

**Supplementary Movie S3 – 3D rendering of a LAMP1 granule.** slncRNA (blue) is clustered at the periphery of an SNA (green)-LAMP1 (red) overlap.

Table 1 – proteins and RNA sequences.

| Protein | NT Sequence | NT Length | AA Sequence | AA Length | Signal Peptide | Notes |
|---|---|---|---|---|---|---|
| ACE2 | ATGTCTAGCTCTAGT | 2220 | MSSSSWLLLSLVAV | 740 | ACE2 | |
| ACE2NB | ATGTCTAGCTCTAGT | 2220 | MSSSSWLLLSLVAV | 740 | ACE2 | D355A |
| delACE2 | ATGTCTAGCTCTAGT | 1887 | MSSSSWLLLSLVAV | 629 | ACE2 | del616-726 |
| LAMP1 | ATGGCCGCCCCTG | 1146 | MAAPGSARRPLLLL | 382 | LAMP1 | |
| GYPA | ATGTACGGCAAGAT | 450 | MYGKIIFVLLLSEIVS | 150 | GYPA | |
| Fetuin | ATGAAAAGCCTGGT | 1101 | MKSLVLLLCLAQLW | 367 | Fetuin | |
| NXST1 | ATGAGCTCCTCCAG | 201 | MSSSSWLLLSLVAV | 67 | ACE2 | |
| NXST2 | ATGAGCTCCTCCAG | 201 | MSSSSWLLLSLVAV | 67 | ACE2 | |
| tdNXST1 | ATGAGCTCCTCCAG | 333 | MSSSSWLLLSLVAV | 111 | ACE2 | |
| mCherry | GTGAGCAAGGGCG | 705 | VSKGEEDNMAIIKE | 235 | | |
| tdPCP | CTAGCCTCCAAAAC | 747 | LASKTIVLSVGEATR | 249 | | |
| mCherry upstream linker | CCACCGGTCGCCA | 15 | PPVAT | 5 | | |
| mCherry downstram linker | CCGCCAGTTGCCA | 15 | PPVAT | 5 | | |
| His-tag | CATCATCACCACCA | 18 | HHHHHH | 6 | | |
| RBD | ATGAAGACCATCAT | 666 | MKTIIALSYIFCLVFA | 222 | HA | |
| sfGFP | TCCAAAGGAGAAGA | 714 | SKGEELFTGVVPILV | 238 | | |

| RNA | NT Sequence | NT Length |
|---|---|---|
| 8xPP7 Cassette | GAGAAACGTTTCGAC | 479 |
| 14xPP7-15xMS2 | GAGAAACGTTTCGAC | 1268 |
